# Spatial confinement reshapes the folding of an ion-stabilized DNA with three-way junction

**DOI:** 10.64898/2026.03.15.711775

**Authors:** Xunxun Wang, Ya-Zhou Shi

## Abstract

Within the densely packed cellular milieu, the structural dynamics and stability of complex DNA architectures are governed by an intricate interplay between spatial confinement and ionic environment. Here we employ our coarse-grained model, DNAfold2, to dissect the mechanistic principles underlying how nanoscale confinement reshapes the folding landscape of a DNA with three-way junction across a broad range of ionic conditions. Our simulations demonstrate that spatial confinement acts as a structural selector, preferentially stabilizing compact, well-defined junction topologies while significantly reducing conformational flexibility in response to ionic strength variations. Through rigorous analysis of structural state populations, we reveal that this behavior results from the entropic exclusion of extended intermediates and unfolded states, effectively reshaping the DNA stability under spatial confinement. Notably, spatial confinement induces a fundamental reshape of the unfolding pathway, suppressing heterogeneous intermediate ensembles and enforcing a highly cooperative transition. These findings establish a unified mechanistic framework for understanding the biophysics of nucleic acid in physiologically crowded environments, with significant implications for de novo design of biomolecular nanomaterials and mechanistic regulation of DNA-processing machinery *in vivo*.

## 1. Introduction

DNA can adopt a wide range of higher-order three-dimensional (3D) conformations beyond the canonical double helix, including hairpins, multi-way junctions, and G-quadruplexes [1-3]. These structural motifs play crucial roles in regulating genetic processes and organizing genomic architecture, while also providing powerful frameworks for the design of programmable DNA nanostructures [4-6]. Among them, multi-way junctions stand out for their intricate topology, which enables their roles in molecular recognition, signal transduction, and nanoscale assembly [5,7]. Inside cells, such structures exist within densely packed and spatially restricted environments [8-10]. For instance, within the crowded environment of the nucleus, where confinement can substantially alter folding behavior and thermodynamic stability [11,12]. While it is broadly accepted that confinement favors structural compaction, its detailed effects on the conformational dynamics and energetic landscape of DNA with multi-way junctions remain poorly understood [13,14]. Hence, a mechanistic understanding of how spatial restriction influences the 3D structure and stability of these complex assemblies is key. It will help explain their cellular functions and guide the rational engineering of DNA-based materials under biologically relevant conditions.

The cooperative interplay between spatial confinement and the ionic environment collectively governs the 3D structure, thermal stability, and thermally unfolding pathway of DNA architectures [15,16]. This coupling is particularly critical for complex DNA motifs such as multi-way junctions [7,17,18], whose structural stability relies on a delicate balance between electrostatic repulsion and ion-mediated stabilization, while spatial confinement entropically favors compact tertiary folds [19]. Understanding this cooperative mechanism is essential to reveal the actual functional behavior of DNA *in vivo*, as the cellular milieu-densely crowded and rich in specific ions-differs fundamentally from idealized low-concentration solutions [20,21]. However, a significant experimental gap persists because techniques such as Small-Angle X-ray Scattering [22-25] and single-molecule imaging [26-28] cannot separate the individual and combined effects of molecular crowding and ionic conditions under physiologically relevant environments. This limitation makes it difficult to achieve a mechanistic interpretation of how confined DNA folding is modulated by ions such as Na^+^ and Mg^2+^, underscoring the need for integrated computational and experimental approaches to bridge this critical knowledge gap.

The computational prediction of DNA faces a fundamental challenge: although deep learning architectures like AlphaFold3 [29,30] and template-based methods such as 3dRNA/DNA [31,32] achieve remarkable accuracy in determining static structures, they remain limited in capturing the non-equilibrium dynamics of folding processes under physiological conditions. This represents a critical gap in our understanding, as cellular DNA operates not in isolation but within an environment characterized by macromolecular crowding and strict ionic regulation [33,34]. Coarse-grained (CG) modeling has emerged as the most powerful strategy to bridge this divide, enabling the simulation of biological-timescale folding while maintaining physical fidelity [35-46].

Under the framework of CG DNA modeling, several established models have been developed, each with distinct strengths. Among them, the 3SPN model represents a nucleotide by its phosphate, sugar, and base. It successfully simulates DNA denaturation and renaturation through its base-stacking and base-pairing interactions [35-38]. The oxDNA model uses three collinear sites and additional vectors to define angle-dependent potentials, accurately describing the structural, mechanical, and thermodynamic properties of both single and double-stranded DNA. Its parameters have been fine-tuned to model large DNA nanostructures like DNA origami [39-41]. TIS-DNA employs nucleotide-specific stacking parameters to reproduce sequence-dependent mechanical and thermodynamic behaviors, including force-extension curves and melting temperatures [44]. These approaches have demonstrated particular value in studying environmental effects; for example, oxDNA simulations have revealed how spatial confinement entropically selects for native-like architectures [39]. However, current CG models suffer from two fundamental simplifications in that they represent molecular crowding as inert excluded volume, ignoring crucial chemical and electrostatic interactions, and rely on Debye-Hückel electrostatics, which fails to capture the specific coordination chemistry of divalent cations like Mg^2+^ that is essential for tertiary structure stabilization [47-49]. Although our previously developed three-bead model represents a significant advance through its ability to predict DNA folding in divalent ion solutions solely from sequence, its current formulation does not incorporate spatial confinement [14,50]. Thus, the development of a unified computational framework that simultaneously accounts for geometric confinement and accurate ion effects remains an outstanding challenge, and its resolution is essential for understanding the functional architecture of DNA in its native cellular environment.

In this study, we employed a representative three-way junction DNA (3WJ) [7] as a model system to systematically explore how spatial confinement modulated its 3D structure and stability under a broad range of ionic conditions. Using our established CG DNA 3D structure model DNAfold2 [14], we first predicted the confined 3D structures and stability profiles of 3WJ across a wide spectrum of ionic concentrations. We subsequently quantified the interplay between spatial confinement and salt-dependent structural behavior. Finally, we performed a comprehensive analysis of the effects of confinement on the thermally unfolding pathways of the 3WJ under varying ionic conditions.

## 2. Materials and methods

### 2.1 Simulation framework and model overview

All simulations were conducted using our established CG model, DNAfold2, which has been extensively validated for predicting DNA 3D structures and stabilities in bulk solution [14]. As shown in Fig. 1, the DNAfold2 model represents each nucleotide with three beads, corresponding to its fundamental chemical groups: a phosphate (P) bead, a sugar (C) bead centered at the C4’ atom, and a nucleobase (N) bead located at the N1 (for pyrimidines) or N9 (for purines) position, respectively [14,50]. The accompanying force field captures essential molecular interactions through explicit terms for sequence-dependent base pairing, base stacking, and electrostatic screening. While the core framework remains unchanged, the current study introduces key extensions to address spatial confinement effects, with complete implementation details available in the Supplementary material.

**Fig. 1:**
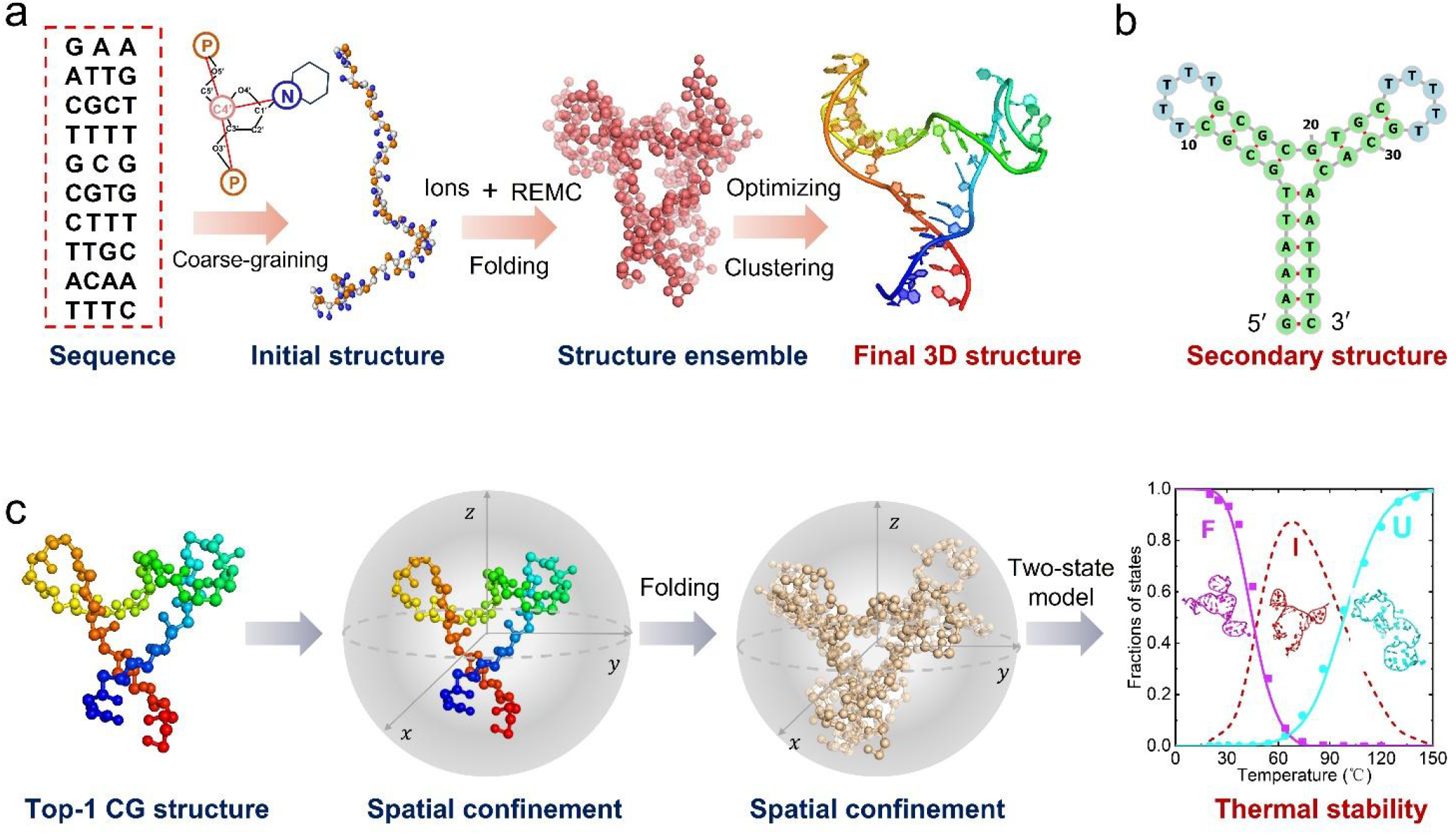
Schematic overview of the CG modeling framework and spatial confinement for the 3WJ. (a) Workflow for predicting the 3D structure from sequence using the DNAfold2 model. The process involves generating an initial CG conformation, conducting replica exchange Monte Carlo (REMC) simulations with ten replicas to sample a diverse ensemble, and selecting a representative top-ranked structure by clustering low-energy conformations. (b) Secondary structure of the DNA three-way junction. (c) Depiction of the confined 3WJ within the spatial constraints modeled by DNAfold2, used for assessing its thermal stability.

### 2.2 Force field with the confinement extension

The total potential energy for a DNA conformation is given by:

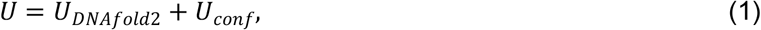

here, the energy function ***U***_***DNAfold*2**_ includes the complete set of energy terms previously developed for the DNAfold2 model [14], integrating both bonded and non-bonded interactions through carefully parameterized energy terms. Bonded potentials (*U*_*b*_, *U*_*b*_, *U*_*b*_) maintain chain connectivity and flexibility, parameterized against high-resolution structural ensembles to ensure geometric accuracy. The non-bonded terms capture essential molecular recognition events: excluded volume (*U*_*exc*_) enforces steric exclusion, while base-pairing (*U*_*bp*_), base-stacking (*U*_*bs*_), and coaxial-stacking (*U*_*cs*_) interactions, which are derived from experimental thermodynamics and optimized through advanced sampling, encode sequence-specific stability. Most significantly, our electrostatic model (*U*_*el*_) represents a key improvement in CG DNA modeling, integrating counterion condensation theory [51] with the tightly bound ion model [52-54] to explicitly capture ion correlation and screening effects that are essential for modeling divalent cation-dependent phenomena. This multi-scale approach enables DNAfold2 to bridge atomic-scale interactions with mesoscale structural transitions, providing significantly improved accuracy in predicting DNA behavior under physiological ionic conditions [54]. Complete parameterization details are provided in the Supplementary material.

A fundamental advancement introduced in this work is the ***U***_***conf***_ term, which provides a rigorous physical treatment of macromolecular crowding through an implicit spatial confinement potential. In environments like the nucleus and macromolecular complexes, where crowding molecules significantly exceed the dimensions of nucleic acid solutes, their collective excluded volume effects can be mathematically mapped onto a geometric confinement model [55,56]. This approach, quantitatively established by Cheung *et al*. for RNA folding thermodynamics [57,58], demonstrates that complex crowding phenomena can be accurately captured through an effective spherical confinement model. Within our DNAfold2 model, the confinement geometry is systematically defined by the size, shape, and packing density of the surrounding macromolecular environment. This theoretical construct enables efficient simulation of crowding-induced structural transitions while avoiding high computational cost of explicit crowder representations. The effective confinement radius follows the established relation [57]:

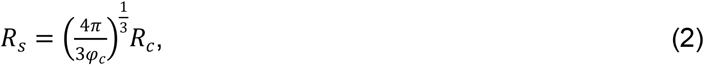

where *R*_*s*_ is the confinement radius, *R*_*c*_ is the crowder radius, and *φ*_*c*_ denotes their volume fraction. Prior theoretical comparisons of various confinement geometries, including planar slits, cylindrical channels, and spherical cavities, revealed that the relative stabilization of compact states is largely geometry-independent [59], thereby justifying the use of spherical confinement as a general and physically grounded representation. Motivated by these insights, we employed spherical cavities of radii *R*_*s*_ =20 Å and 40 Å (computed from eq 2) to investigate crowding-induced modulation of the folding landscape of a DNA three-way junction across a broad range of Na^+^ concentrations (Fig. 1a). To represent the steric boundary, we adopted a purely repulsive hard-wall potential-shown to be effectively equivalent to softer repulsive forms such as 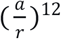 [60] for confined biomolecules-defined as [61]:

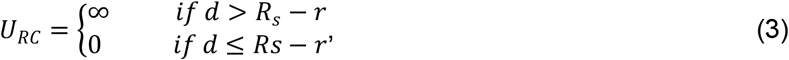

where *d* is the distance from the bead to the center of confinement and *r* is its van der Waals radius.

### 2.3 Simulation and analysis protocols

We employed the Replica Exchange Monte Carlo (REMC) method to achieve adequate sampling of the conformational landscape under both confined and bulk conditions [54,62,63]. Our temperature set consisted of ten replicas, ranging from 25 °C to 110 °C, ensuring efficient traversal over folding barriers. Conformational sampling within each replica was driven by pivot moves following to the standard Metropolis criterion [64]. Notably, in our DNAfold2 model, exchanges between adjacent temperatures were periodically proposed and accepted with a probability of *p* = *min* (1, *exp* (− Δ)), where Δ = *β*_*j*_*E*(*x*_*i*_, *T*_*j*_) + *β*_*i*_*E*(*x*_*j*_, *T*_*i*_)− *β*_*i*_*E*(*x*_*i*_, *T*_*i*_)− *β*_*i*_*E*(*x*_*j*_, *T*_*j*_)), and *E*(*x*) is the potential energy of conformation ***x***. This generalized acceptance criterion ensures the preservation of detailed balance even for systems with explicit temperature-dependent interactions [65,66].

### 2.4 Thermal stability and thermally unfolding pathway analysis

The thermodynamic properties of the 3WJ were extracted from the REMC trajectories using the Weighted Histogram Analysis Method (WHAM) [67-69]. To quantitatively track unfolding pathway, we defined a set of discrete structural states (e.g., fully folded, unfolded, and specific intermediate states) based on the presence or absence of native helical stems (Fig. S1). The population of each state as a function of temperature was calculated, allowing us to derive melting temperatures (*T*_*m*_) and characterize the thermally unfolding pathways.

### 2.5 Selection of top-scoring coarse-grained 3D Structures

To identify the predominant low-energy conformations, we extracted the 1,000 lowest-energy structures from the 25°C replica ensemble [43]. These structures were subsequently subjected to RMSD-based hierarchical clustering analysis. The clustering procedure iteratively identified the largest cluster within a predefined cutoff, removed it, and repeated the process until all conformations were assigned. The centroid of the most populous cluster-defined using a normalized threshold of 0.1 Å per nucleotide (equivalent to 7 Å for the 70-nt sequence)-was selected as the representative structure [70]. Finally, the medoids of the three largest clusters and the lowest-energy decoy conformation were selected as initial structural predictions.

### 2.6 Structure prediction and reconstruction

To obtain high-fidelity all-atom structures, we refined selected CG structures through a multi-stage computational pipeline. First, Monte Carlo optimization at room temperature (25 °C) enhanced structural accuracy using differentiated potentials for base-paired (Para_helix_) and unpaired (Para_loop_) regions. These refined CG structures were then converted to all-atom representations through a fragment-based reconstruction algorithm. We employed a pre-assembled library of nucleotide and base-pair conformers, iteratively aligning each CG unit to sequence-matched fragments and selecting optimal matches through RMSD minimization (Fig. S2). Finally, we performed energy minimization with geometric restraints using QRNAS [71] to resolve steric clashes and ensure proper stereochemistry, yielding physically realistic structures for subsequent analysis.

### 2.7 The DNAfold2 computational framework

Figure 1 presents the multi-step workflow of DNAfold2 model for predicting DNA 3D structures from sequence. First, an initial conformation is generated using bonded (*U*_*bond*_) and excluded-volume (*U*_*exc*_) potentials (eq 1). This structure subsequently undergoes REMC sampling with Para_loop_-derived bonded potentials to model intrinsic chain flexibility. Finally, low-energy structures are selected from the lowest-temperature replica (e.g., 25°C) through clustering analysis [72]; (4) To improve geometric fidelity, we refined the top-scoring conformations using Monte Carlo simulations at 25°C. The refinement employed structure-specific parameterization: Para_helix_ potentials enforced canonical geometry in base-paired stems, while Para_loop_ potentials preserved the conformational flexibility of loop regions [73]; (5) Finally, the refined CG structures were converted into final all-atom structures through all-atom rebuilding algorithm from DNAfold2 model.

To evaluate the performance of the DNAfold2 model, we employed two principal metrics: root-mean-square deviation (RMSD) and F1-score. The RMSD quantifies global structural deviations between predicted and native structures, with the top-1 RMSD specifically indicating the accuracy of the highest-ranked prediction [74,75]. The F1-score assesses the local base-pairing fidelity by comparing predicted contacts with the native pairing pattern of structure [76,77]. The F1-score is calculated by *F*1 = 2 × *PR* × *SN*/(*PR* + *SN*), where precision (PR) is defined as *PR* = *TP* /(*TP* + *FP*) and sensitivity (SN) as *SN* = *TP*/(*TP* + *FN*), with TP, FP, and FN denoting true positives, false positives, and false negatives, respectively. Furthermore, predictive reliability for DNA thermodynamic stability was defined by the deviation between predicted and experimental melting temperatures (*T*_*m*_), with smaller deviations indicating higher accuracy in predicting thermal denaturation behavior.

## 3. Results and discussion

Employing our DNAfold2 model, we quantitatively dissect how spatial confinement governs the architecture and energetics of the 3WJ, which is a fundamental structural motif whose behavior under physiological crowding remains poorly characterized [7]. Through systematic sampling across physiologically relevant regimes of confinement geometry, ionic environment, and thermal fluctuations, we establish a quantitative relationship between nanoscale spatial constraints and the thermodynamic stability of three-way junction DNA. Our multi-parametric approach reveals how geometric confinement restructures the folding landscape and redefines the ionic dependence of complex DNA architectures under conditions mimicking the crowded cellular environment.

### 3.1 Validation of DNA 3D structure with existing methods

To evaluate the predictive accuracy of our DNAfold2 model for 3WJ taking a 37-nt DNA (the 3WJ with sequence of 5’-GAAATTGCGCTTTTTGCGCGTGCTTTTTGCACAATTTC-3’) as an example without an available experimental 3D structure (Fig. 2) [7], we compared its predicted structure against those generated by two representative methods: 3dRNA/DNA [31] and AlphaFold3 [29]. All predictions were generated using sequence information as the sole input. Furthermore, to enable a comprehensive comparison with 3dRNA/DNA, we conducted additional predictions using its structure-guided mode by inputting the native secondary structure along with the sequence. All 3D structures were retrieved from the respective official web servers under default parameter settings.

**Fig. 2:**
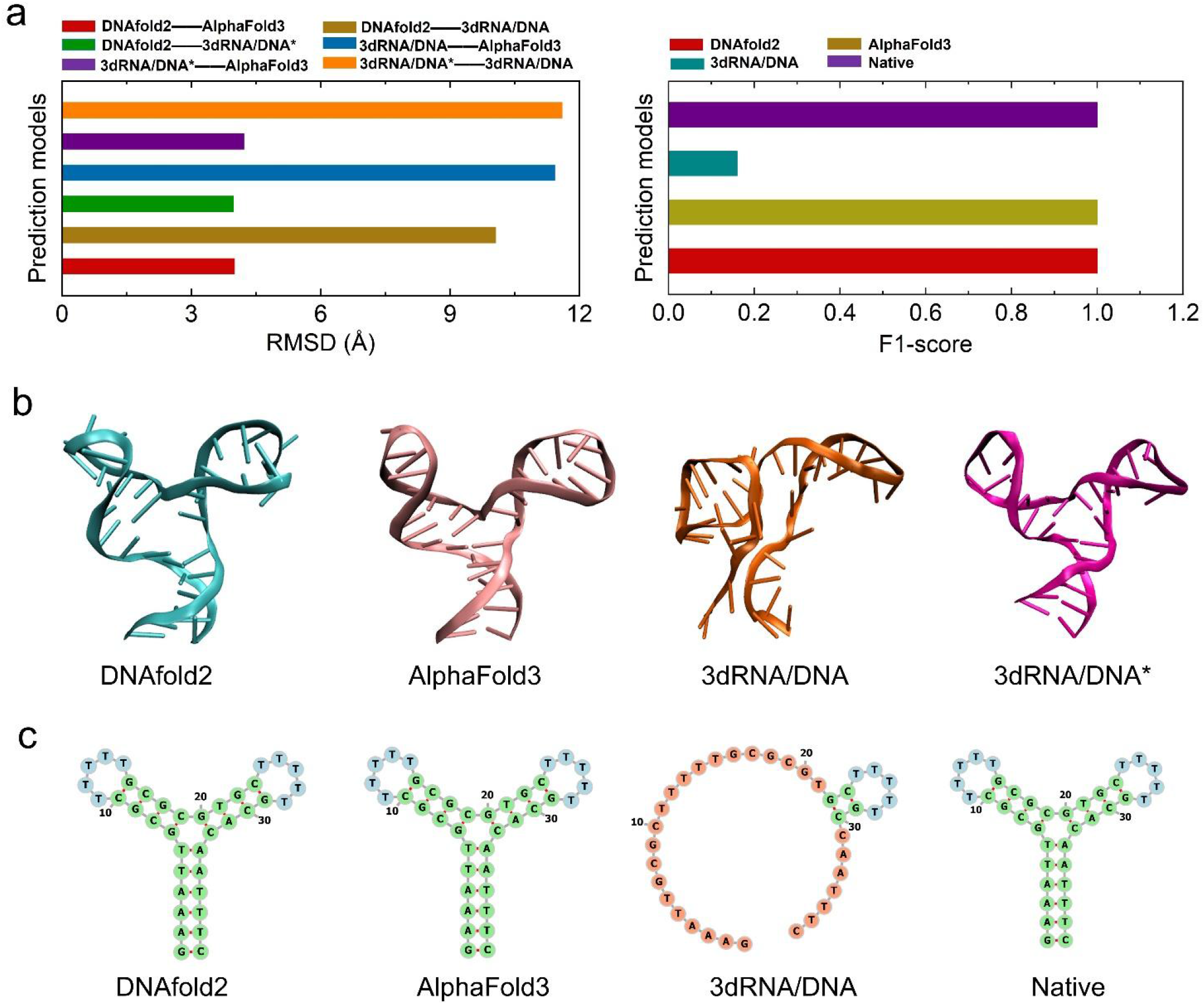
Comparison of predicted secondary structures and 3D structures of the 3WJ using three DNA structure prediction models . (a) Quantitative evaluation of top-1 RMSD (left) and F1-socre (right) predicted 3D structures of the 3WJ. (b) Visual comparison of predicted 3D structures of DNA by different models: DNAfold2 (cyan), AlphaFold3 (pink), 3dRNA/DNA (orange), and 3dRNA/DNA* (magenta). (c) Comparison of predicted secondary structures of DNA generated by DNAfold2, AlphaFold3, and 3dRNA/DNA models. Here, 3dRNA/DNA predicts structures based on DNA sequences, while 3dRNA/DNA* represents a modified version that predicts structures based on Native secondary structures. The secondary structures and 3D structures of the DNA were visualized using the ViennaRNA software [83] and the VMD (Visual Molecular Dynamics) software [84], respectively.

Since the experimental structure was unavailable, we evaluated prediction accuracy by assessing the structural consensus among models. As shown in Fig. 2, the prediction of DNAfold2 forms a tight cluster with two high-accuracy benchmarks: the structure-guided 3dRNA/DNA* and AlphaFold3. Notably, DNAfold2 demonstrates nearly identical agreement with both, yielding RMSD values of 3.9 Å to 3dRNA/DNA* and 4.0 Å to AlphaFold3, while the mutual RMSD between AlphaFold3 and 3dRNA/DNA* is also comparable at 4.1 Å (Fig. 2). Here, 3dRNA/DNA* refers to a method that predicts 3D structures from secondary structure. This strong consensus clearly contrasts with the sequence-only 3dRNA/DNA prediction (RMSD >10.0 Å), which emerges as a clear outlier. As shown in Fig. 2, independent validation is provided by secondary structure analysis: both DNAfold2 and AlphaFold3 achieve a perfect F1-score of 1.0 against the experimental secondary structure, compared to only 0.15 for 3dRNA/DNA. This convergence with two independent and accurate methods, at both the 3D and secondary structure levels (Figs. 2b and c), provides compelling evidence that DNAfold2 has captured the correct native structure, underscoring the critical role of accurate secondary structure information for successful 3D modeling.

In summary, the convergent evidence from 3D structural consensus and perfect secondary structure recovery provides robust, dual-validation that the DNAfold2 model has accurately predicted the native 3D structure of the 3WJ. This highly precise prediction not only validates the reliability of our computational approach but also establishes a solid foundation for investigating its folding under spatial confinement.

### 3.2 Spatial confinement compacts unfolded states and attenuates ionic modulation

Our findings establish that spatial confinement acts as a conformational editor, selectively reshaping the structural landscape of the 3WJ in a manner that depends on its folding state. As shown in Fig. 3, we systematically investigated the 3D structure of the 3WJ under varying monovalent ion concentrations, temperatures, and degrees of spatial confinement (confinement radii *R*_*s*_ of 20 Å and 40 Å). Figs. 3a-f represent the resulting radius of gyration (*R*_*g*_) and number of predicted base pairs (*N*_*bp*_). The response to confinement (*R*_*s*_ =20 Å) is highly asymmetric when compared to the unconfined state (*R*_*s*_ =∞). At 25 °C, the natively folded junction (*R*_*g*_ ≈15-16 Å) shows minimal compaction (Δ*R*_*g*_ ≈2-3 Å). In contrast, at 110 °C, the unfolded ensemble undergoes dramatic restructuring, with its *R*_*g*_ reduced by ∼14 Å relative to its unconfined dimensions. This significant asymmetry reveals a fundamental principle: spatial confinement (***R***_*s*_ =20 Å) acts not by distorting energetically favored native states, but by eliminating the large-scale conformational fluctuations unique to denatured states. The physical basis for this selective editing lies in the intrinsic extensibility of the junction. As shown in Fig. 4, we quantified this extensibility by *R*_*max*_, the maximum distance between any CG bead and the conformational centroid. Under non-confinement conditions (*R*_*s*_ = ∞), the unfolded ensemble samples highly extended conformations (*R*_*max*_ ≈50 Å) that exceed the spatial constraints imposed by confinement, while the folded state naturally resides within these geometric bounds (*R*_*max*_ ≈32 Å).

**Fig. 3:**
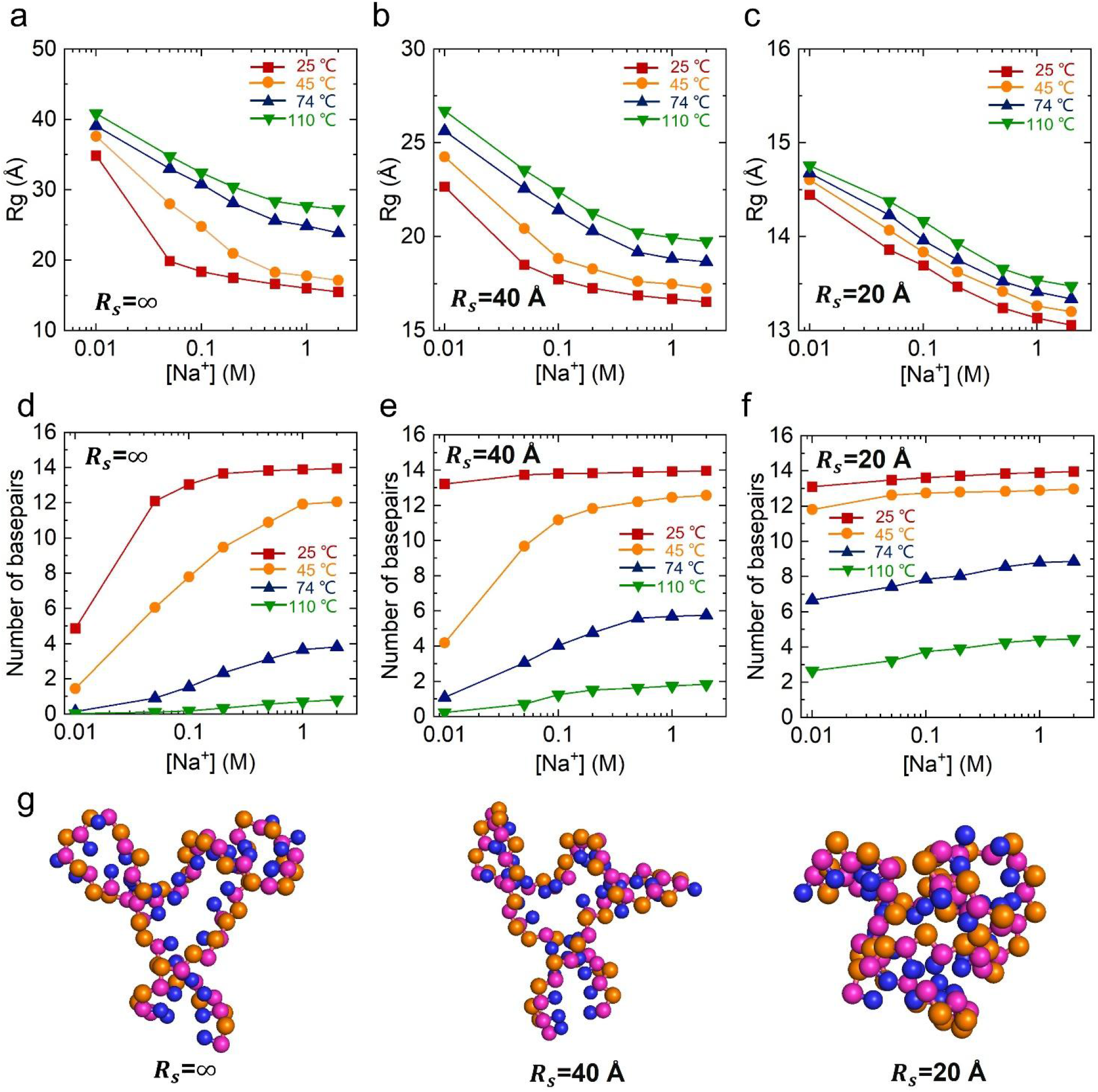
The influence of spatial confinement on the salt dependence of the 3WJ. (a-c) Radii of gyration (*R*_*g*_) as a function of Na^+^ concentration for 3WJ at different temperatures and confinement conditions: (a) *R*_*s*_=∞, (b) *R*_*s*_ = 40 Å, and (c) *R*_*s*_ = 20 Å. (d-f) Fractions of native base pairs formed as a function of Na^+^ concentration under different temperatures and confinement conditions: (d) *R*_*s*_ =∞, (e) *R*_*s*_ = 40 Å, and (f) *R*_*s*_ = 20 Å. (g) The 3D structures predicted by DNAfold2 under different confinement radii.

**Fig. 4:**
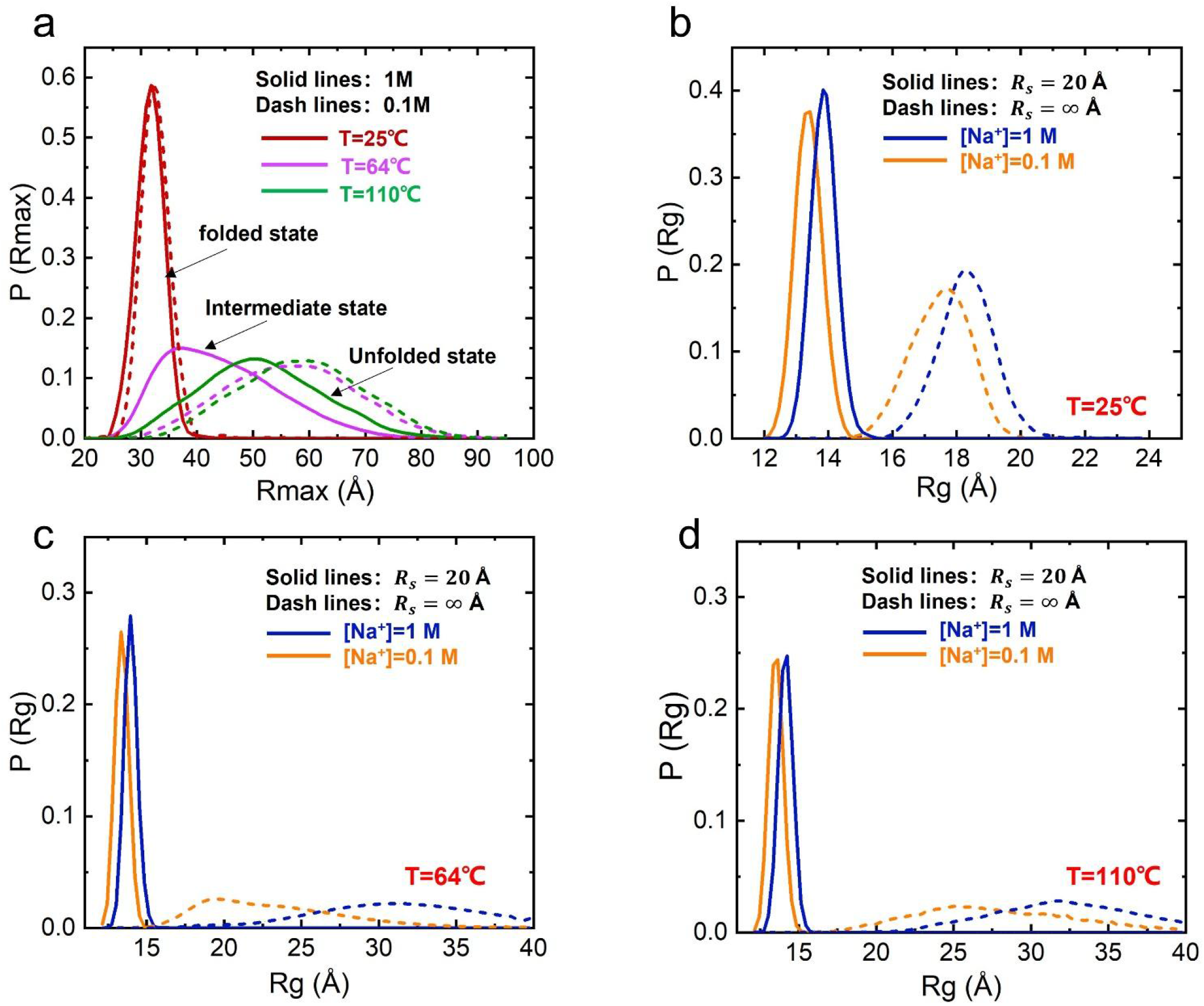
The structural state distributions of the 3WJ under different degrees spatial confinement. (a) Distributions of the maximum extension (*R*_*max*_) for the 3WJ at 1 M [Na^+^] across different temperatures. (b-d) Distributions of the radius of gyration (*R*_*g*_) for the 3WJ under varying [Na^+^] and temperatures: (b) 25 °C, (C) 64 °C, and (D) 110 °C. In (b-d), solid and dashed lines correspond to confined (*R*_*s*_ = 20 Å) and unconfined (*R*_*s*_ =∞) conditions, respectively.

Furthermore, our analysis reveals that geometric control directly modulates the coupling between ion atmosphere and DNA conformation. As shown in Figs. 3b-c, in the absence of spatial confinement, elevating the Na^+^ concentration from 0.1 M to 1 M compacts the unfolded DNA ensemble by Δ*R*_*g*_ ≈7 Å, a consequence of enhanced electrostatic screening. However, under strong spatial confinement (*R*_*s*_ =20 Å), this pronounced ionic response is nearly abolished, with a markedly reduced compaction of only Δ*R*_*g*_ ≈1 Å. This stark contrast clearly demonstrates that physical confinement can dominate electrostatic influences. Mechanistically, both confinement and ions operate on the same degree of freedom-molecular extension. Once the available spatial volume is saturated by confinement-induced compaction, further electrostatic screening cannot drive additional compaction, effectively decoupling structural behavior from ionic environment. This ‘conformational saturation’ mechanism, wherein geometric constraints exhaust the available spatial degrees of freedom, explains the attenuation of ionic modulation.

The thermodynamic effects of this conformational editing are directly reflected in base-pairing stability. Our analysis reveals that confinement significantly enhances native contact formation specifically under intermediate ionic conditions, where the junction retains partial structural plasticity, by destabilizing extended, high-entropy intermediates and shifting the equilibrium toward compact, ordered states. In contrast, confinement exerts minimal influence when the 3WJ is either fully folded under stabilizing conditions of high salt (1 M Na^+^) and low temperature (25 °C), or completely denatured under challenging conditions of low salt (0.01 M Na^+^) and high temperature (110 °C), as quantified by the predicted number of base-pairs (*N*_*bp*_); see Figs. 3d-f. This clear contrast-from a near-maximum *N*_*bp*_ of ∼14 in the native state to an *N*_*bp*_ of ∼0 in the unfolded state-definitively reveals that the primary role of confinement is not as a universal stabilizer, but as a selective enhancer of folding cooperativity.

Additionally, the progressive DNA compaction under increasing spatial confinement, revealed in Fig. 3g, is not merely a physical phenomenon but has direct implications for gene regulation. This entropy-driven compaction which is forced by the exclusion of extended conformations suggests a physical mechanism for the control of DNA accessibility. In the crowded nuclear environment, such confinement-induced compaction could naturally modulate the exposure of specific gene regions to transcription machinery, thereby influencing gene expression patterns without direct chemical modification.

In summary, our findings establish a unified biophysical mechanism through which spatial confinement governs nucleic acid architecture: by entropically excluding extended states, it selectively (*i*) compacts unfolded ensembles, (*ii*) attenuates salt-dependent structural responses, and (*iii*) stabilizes native base pairing under intermediate folding conditions. This mechanistic framework extends beyond the specific case of the 3WJ, offering a general principle for understanding nucleic acid behavior in geometrically constrained environments.

### 3.3 Confinement enhances thermal stability by entropic exclusion and alters salt dependence

The functionality of DNA is inherently linked to both its 3D structure and thermal stability [78-80]. As show in Fig. 5, beyond 3D structure prediction, we applied DNAfold2 to quantify the thermal stability of the 3WJ under varying monovalent ion concentrations and degrees of spatial confinement. Using REMC simulations across a wide temperature range (Figs. 5a-c), we classified equilibrium conformations into three dominant states-fully folded (F), intermediate (I), and unfolded (U)-based on native secondary structure features (Figs. 5d-f). As shown in Figs. 5g-i, fitting the temperature-dependent populations to a two-step melting model yielded *T*_*m*1_ and *T*_*m*2_ for the F → I and I → U transitions, respectively [73]. Validation against experimental data at low salt ([Na^+^] = 0.1-0.2 M) in the absence of confinement (*R*_*s*_ =∞) showed deviations below 3 °C, confirming the predictive reliability of our DNAfold2 model; see Tables S4-S6. Thus, this validated model provides a robust foundation for investigating the coupled effects of ionic environment and spatial confinement.

**Fig. 5:**
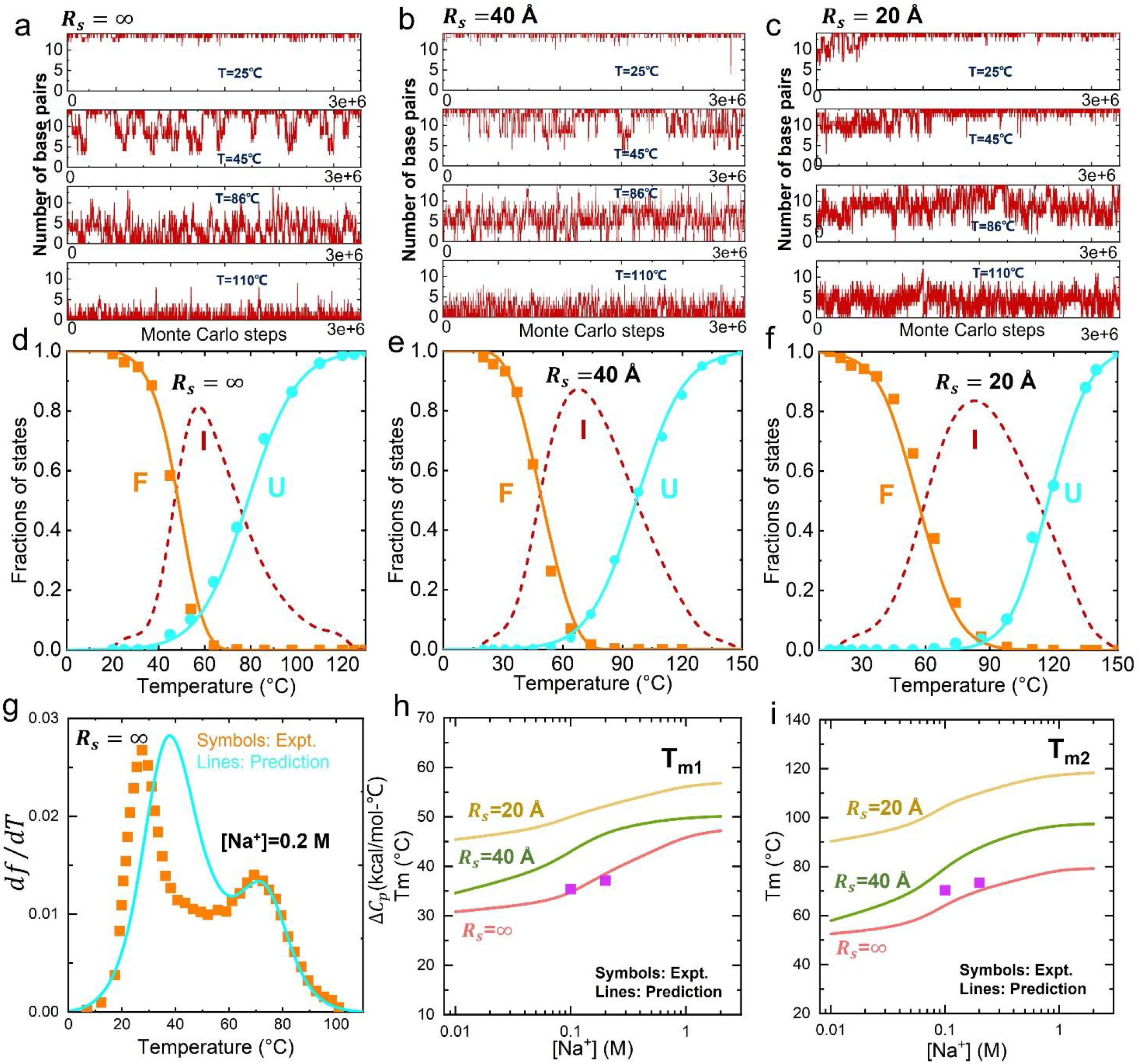
Effect of spatial confinement on the thermal stability of the 3WJ. (A-C) Time evolution of the base-pair fraction in 3WJ at different temperatures (110°C, 86°C, 45°C, and 25°C, from bottom to top) under varying degrees of spatial confinement: (a) *R*_*s*_ = ∞ (non-confinement), (b) *R*_*s*_ = 40 Å, and (c) *R*_*s*_ = 20 Å. (d-f) Temperature-dependent fractions of the folded (F, orange), Intermediate (I, red), and unfolded (U, cyan) states of 3WJ in 1M NaCl under confinement: (d) *R*_*s*_ = ∞, (e) *R*_*s*_ = 40 Å, and (f) *R*_*s*_ = 20 Å. (g) Comparison between experimental melting data (orange symbols) and DNAfold2 predictions (cyan curves). (h, i) Predicted melting temperatures *T*_*m*1_ (h) and *T*_*m*2_ (i) as functions of the spatial confinement radii *R*_*s*_s, from DNAfold2 model.

Both increasing ionic strength and spatial confinement enhance 3WJ stability, but via distinct physical mechanisms. As shown in Figs. 5h and i, under unconfined conditions (*R*_*s*_ =∞), raising [Na^+^] from 0.01 M to 2 M increases *T*_*m*1_ and *T*_*m*2_ by ∼16°C and ∼28°C, reflecting electrostatic screening of backbone repulsion. The effect of confinement is starkly different at high salt. Under strong spatial confinement (*R*_*s*_ =20 Å) and at 2 M [Na^+^], *T*_*m*1_ increases to a value that is ∼9.6 °C higher than its unconfined counterpart, while *T*_*m*2_ increases to a value that is ∼39 °C higher. Mechanistically, confinement primarily restricts the entropic freedom of extended conformations, strongly destabilizing the unfolded state. Consistently, analysis of *R*_*max*_ distributions shows that the U state is substantially more extended than both F and I states (see Fig. S3); confinement compresses the U ensemble dramatically, effectively stabilizing intermediate and folded states by reducing accessible high-entropy configurations.

This entropic-limiting mechanism also explains why confinement attenuates salt-dependent stabilization. By restricting the spatial fluctuations of extended states, confinement reduces the structural and electrostatic contrast between folding states. As a result, the incremental benefit of ionic screening is reduced, with the result that the differences in melting temperatures between low and high salt conditions decrease under confinement, particularly for the I→U transition, where the U state converges toward the I state in terms of compactness and effective charge density. This decoupling of structural stability from ionic environment may have critical biological implications. Within the nucleus, where molecular crowding creates naturally confined spaces, our findings suggest a conformational buffering effect. Such geometric buffering could ensure the stability of essential DNA architectures, like replication forks or transcription bubbles, against local and transient fluctuations in ion concentration, thereby promoting genomic stability. In essence, confinement shifts the folding landscape from one dominated by electrostatic modulation to one constrained predominantly by geometry and conformational entropy.

Thus, these findings support a clear physical picture: spatial confinement stabilizes 3WJ primarily by restricting the conformational entropy of extended ensembles, which (*i*) significantly compacts the unfolded state, (*ii*) enhances intermediate-state stability relative to the fully folded state, and (*iii*) weakens the dependence of folding transitions on ionic strength. This unified mechanism aligns with observations in crowded RNA systems, providing a generalizable framework for understanding how cellular confinement and macromolecular crowding modulate nucleic-acid folding landscapes [13] .

### 3.4 Confinement reshapes the unfolding pathway by selecting compact Intermediates

Given the critical role of unfolding pathways in the structural transitions of 3WJ [7,81,82], we further performed a detailed analysis under various conditions to assess the effect of spatial confinement on the unfolding process. Based on our simulations at defined temperature and salt conditions, the intermediate state (I) can be further categorized into six distinct structural states: intermediate states I1 (Stem 1 melted), I1’ (Stem 3 melted), I1’’ (Stem 2 melted), I2 (Stems 1 and 3 melted), I2’ (Stems 1 and 2 melted) and I2’’ (Stems 2 and 3 melted), respectively; see Fig. S1 in the Supplementary material. The population fractions of these structural states were computed across different salt concentrations and levels of spatial confinement.

As shown in Figs. 6a (state populations) and b (unfolding pathways), in the absence of spatial confinement (*R*_*s*_ =∞ Å), at 1 M [Na^+^]. As the temperature increases from ∼30°C to ∼50°C, the fraction of F decreases from ∼96% to ∼38%, while the fractions of intermediate states I1, I2, and I2’ gradually increase to ∼37%, ∼20%, and ∼4%, respectively. Between ∼50°C and ∼70°C, the fractions of the F and I1 further decrease to ∼1% and ∼14%, whereas I2 and I2’ reach their peak values of ∼51% and ∼8%, respectively. At ∼110°C, the system transitions into the unfolded state (U). These observations suggest that the dominant unfolding pathway of the 3WJ follows F → I1 → I2 → U, with I2 being the most populated intermediate at ∼70°C. Additionally, two minor pathways are identified: F→I2→U and F→I1→I2’ →U, with the former having a significantly higher flux than the latter. In the spatial confinement *R*_*s*_ = 40 Å (20 Å), at 1 M [Na^+^]. As the temperature increases from ∼30°C to ∼60°C, the fraction of F decreases from ∼99% (∼99%) to ∼12% (∼49%), while the fractions of intermediate states I1, I1’, I2, and I2’ gradually increase to ∼59% (∼38%), ∼2% (∼5%), ∼19% (∼3%), and ∼4% (∼2%), respectively. Between ∼60°C and ∼80°C, the fractions of the F and I1 further decline to ∼0% (∼9%) and ∼25% (∼50%), whereas I2 and I2’ increase to ∼44% (∼21%) and ∼9% (∼7%), respectively. At ∼110°C, the system transitions into the unfolded state (U). These observations indicate that the primary unfolding pathway of the 3WJ follows the sequence F→I1→I2→U, in which the intermediate state I2 is the most abundant at temperatures of approximately 80-100°C. Additionally, two minor pathways are identified: F→I1’ →I2→U and F→I1→I2’ →U, with the former having a slightly higher flux than the latter. The above results indicate that spatial confinement can alter the unfolding pathway of the 3WJ. Spatial confinement effectively modulates the unfolding pathway of the molecule by selectively altering the stability and distribution of intermediate states. Specifically, under confined conditions, a more compact intermediate state, I1’, emerges along the pathway (Figs. 6a and b). The reason for this is that I1 ‘ exhibits a smaller average radius of gyration (*R*_*g*_) and maximum distance *R*_*max*_ compared to the I1 intermediate; see Fig. 6c. As a result, spatial restriction significantly suppresses the formation of the I1 state, thereby stabilizing the otherwise unstable I1’ state. Furthermore, as the confinement size (*R*_*s*_) decreases, the probability of the I1’ state gradually increases (Fig. 7a). This indicates that stronger spatial confinement more effectively suppresses the I1 state, thus preferentially favoring the population of the I1’ state. These findings reveal that spatial confinement alters the unfolding pathway by entropically and geometrically selecting for compact conformations that fit the confined space. The suppression of the more extended I1 state and the concurrent stabilization of the compact I1’ intermediate directly reshape the underlying free-energy landscape, effectively “reprogramming” the dominant flux through the available pathways.

**Fig. 6:**
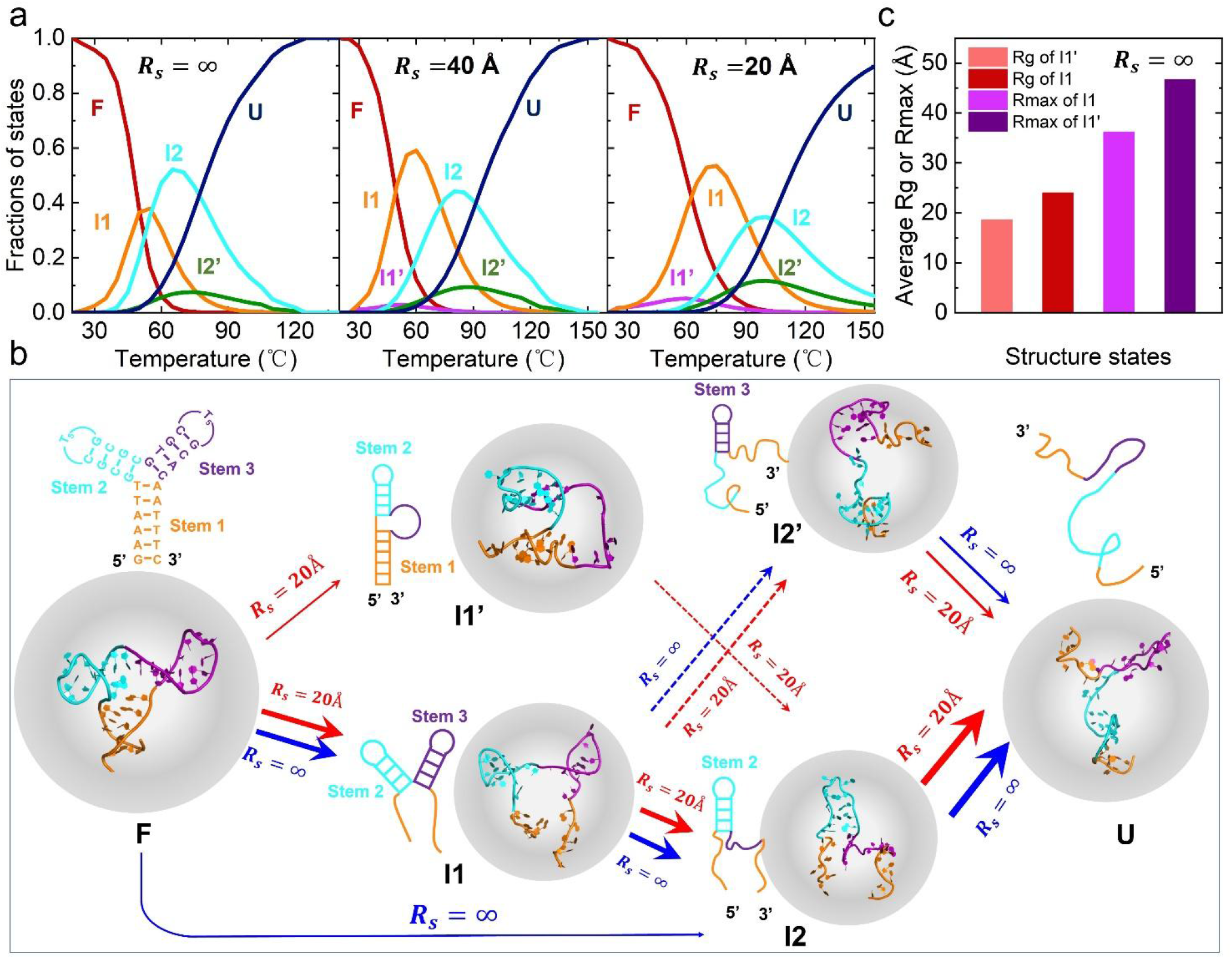
Predicted thermally unfolding pathway of the 3WJ at 1 M [Na^+^] with/without spatial confinement using DNAfold2 model. (a) Temperature-dependent distributions of structural states during the unfolding of 3WJ, showing the fractions of fully folded (F), intermediate (I1, I1’ I2, I2’), and fully unfolded (U) states; *R*_*s*_ = ∞ (non-confinement, left panel), *R*_*s*_ = 40 Å (medium panel), and *R*_*s*_ = 20 Å (right panel). Here, F represents the fully folded state, I1 corresponds to the intermediate state with Stem 1 and 3 melted, I2 and I2’ represent the hairpin intermediates with Stem 2 and Stem 3 retained, respectively, and U denotes the fully unfolded state. (b) Schematic representation of the structure transitions along the unfolding pathway inferred from the state fractions shown in (a). (c) The average *R*_*g*_ and *R*_*max*_ of I1 and I1’ states.

**Fig. 7:**
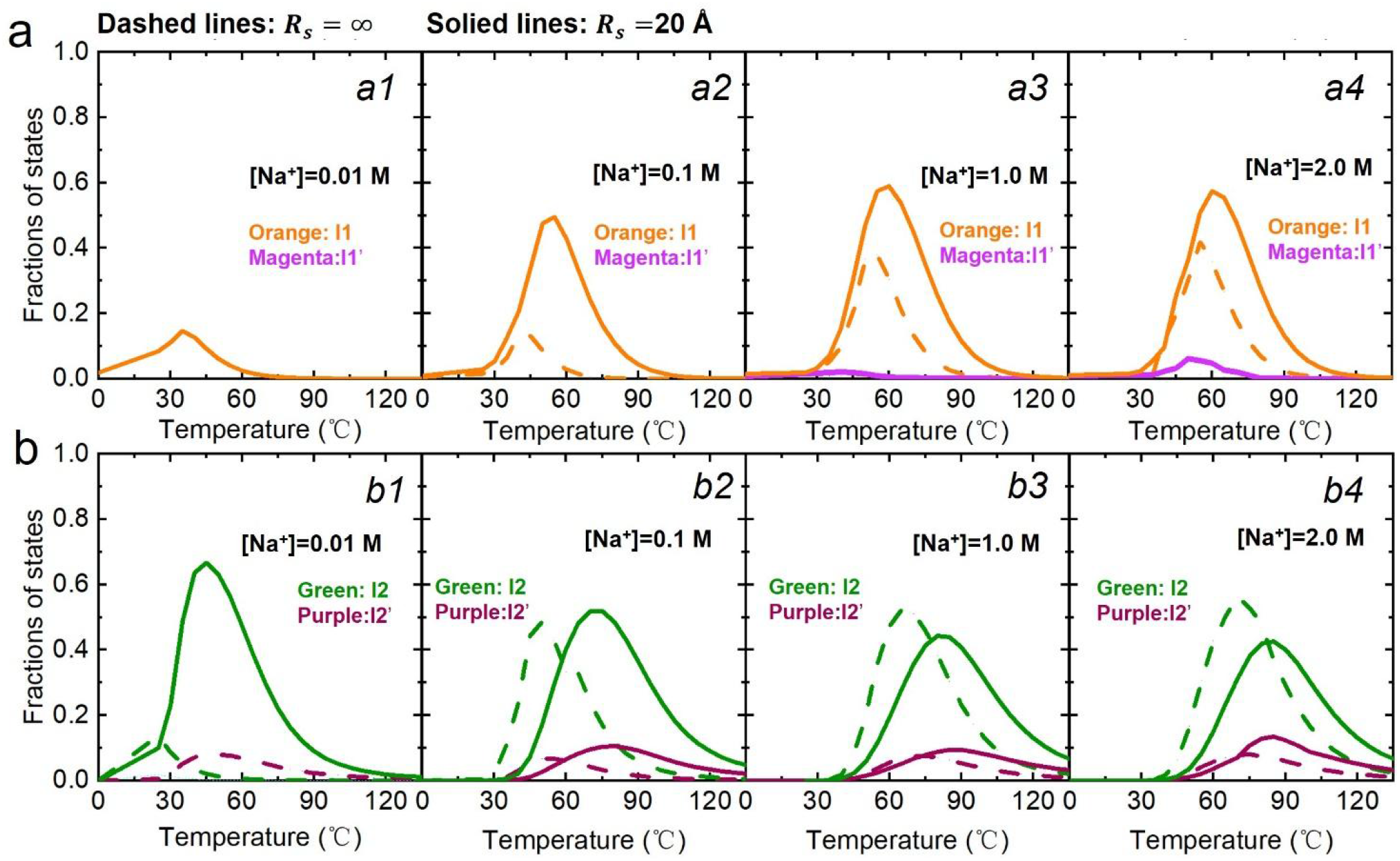
Temperature-dependent fractions of the I1, I1’, I2, and I2’ states during thermally unfolding of the 3WJ under unconfined (*R*_*s*_ = ∞, dashed lines) and confined (*R*_*s*_ = 40 Å, solid lines) conditions. (a) Fractions of the I1 and I1’ intermediate states at (*b1*) 0.01 M, (*b2*) 0.1 M, (*b3*) 1 M, and (*b4*) 2 M [Na^+^]. (b) Fractions of the I2 and I2’ intermediate states at (*c1*) 0.01 M, (*c2*) 0.1 M, (*c3*) 1 M, and (*c4*) 2 M [Na^+^].

Furthermore, as shown in Fig. 7, spatial confinement under high salt conditions significantly alters the unfolding pathway of the 3WJ. For instance, at 1 M [Na^+^], introducing spatial confinement (*R*_*s*_ = 20 Å) shifts the maximum population fractions of the I2 and I2 ‘ states from ∼51% and ∼8% to ∼35% and ∼12%, respectively. This redistribution arises from the difference in conformational dimensions between the two intermediates, as the *R*_*max*_ distribution of the I2 state peaks at ∼50 Å, whereas the more compact I2 ‘ state peaks at ∼33 Å; see Fig. S4 in the Supplementary material. Consequently, spatial confinement more strongly suppresses the extended I2 conformation, reducing its population while favoring the compact I2’ state. Notably, under confinement, increasing the ion concentration from 0.1 M to 2 M further suppresses the I2 state and enriches the I2’ state (Fig. 7b), a trend opposite to that observed in the unconfined system. This concentration-dependent effect arises because the I2 state, with its intrinsically larger *R*_*g*_ and *R*_*max*_, becomes even more expanded at higher salt due to enhanced electrostatic screening, thereby experiencing stronger entropic suppression under spatial restriction and accelerating the shift toward the I2 ‘ state. This phenomenon can be attributed to the strong intramolecular electrostatic repulsion under low-salt conditions, which balances the compression effect induced by spatial confinement.

In conclusion, our study demonstrates that spatial confinement functions as a powerful steric filter, fundamentally reshaping the unfolding pathway of the 3WJ. The core mechanism is an entropic selection process, finely tuned by both physical constraint and electrostatic environment. Under high ionic strength, this process drives spatial confinement to impose a severe entropic penalty on expanded conformations, thereby selectively stabilizing compact intermediates (I1 ‘, I2 ‘) over their extended counterparts (I1, I2); see Fig. 7.

More broadly, these findings extend beyond a single DNA structure to illustrate a general physical principle for cellular biomolecular organization. The crowded and confined intracellular milieu is not a passive background but an active regulatory element that can steer conformational transitions through the interplay between spatial entropy and electrostatic screening. This principle implies that the functional folding pathways observed *in vitro* may not represent the full spectrum *in vivo*, but may be narrowed by the native cellular environment, which actively suppresses certain pathways while enhancing others to ensure efficient and correct folding. Consequently, our work provides a mechanistic framework for understanding how nanoconfinement in organelles, chaperone cavities, or the ribosome exit tunnel can contribute to protein and DNA homeostasis. From a technological perspective, these insights offer a guide for biomimetic control in nanotechnology, suggesting that artificial confinement can be rationally designed to manipulate the assembly pathways of nucleic acid nanostructures or to modulate the stability of therapeutic DNAs.

## 4. Conclusion

In the cellular environment, DNA folding is governed by the complex interplay of electrostatic forces and spatial constraints. By integrating a spatially confining potential into our coarse-grained model, we have decoupled these effects to reveal that molecular crowding fundamentally reshapes the folding landscape of a DNA with three-way junction (3WJ), not merely by stabilizing its native state, but by actively editing its ensemble of unfolding pathways.

Our findings reveal that spatial confinement functions as a dominant entropic editor of DNA stability and folding pathways. It exerts this role by imposing a severe entropic penalty on extended conformations, which leads to two profound consequences. First, it rewrites the rules of electrostatic sensitivity by significantly attenuating the well-established salt-dependence of DNA stability under confinement. This occurs because the physical wall preemptively excludes the very expanded states that would otherwise be stabilized by electrostatic screening at high ionic strength. Second, this entropic exclusion dramatically elevates thermal stability, effectively raising the free-energy barrier for the terminal unfolding step from intermediates to the fully unfolded state.

Most significantly, our results demonstrate that the role of confinement extends beyond passive stabilization to active pathway editing. Under high-salt conditions, it selectively stabilizes a compact intermediate (I1 ‘), thereby altering the unfolding pathway. This positions spatial confinement not as a mere background stabilizer, but as an allosteric regulator that acts at a distance, steering conformational transitions by reshaping the underlying free-energy landscape.

In essence, this work reveals that in a crowded milieu, the folding pathway of a DNA structure is not solely determined by its sequence or modulated by ions, but is decisively authored by the physical confines of its environment. This represents a fundamental change in perspective, which moves from viewing confinement as a passive constraint to recognizing it as an active editor, thereby providing a new conceptual framework for understanding nucleic acid topology and function in the spatially organized, crowded interior of the cell.

## Acknowledgements

We are grateful to Profs. Yang Yu (Guizhou Medical University), Qiude Li (Guizhou Medical University), Zhi-Jie Tan (Wuhan University), and Bengong Zhang (Wuhan Textile University) for valuable discussions, and we would like to acknowledge computing resources from the Super Computing Center of Guizhou Medical University. This work supported by the grants from Guizhou Provincial Science and Technology Program (No. TR25021), Guizhou Medical University High-Level Talent Scientific Research Startup Fund (No. 26242020163), and the National Science Foundation of China (No. 32571442).

## References

[1] Z. D. Smith, S. Hetzel, A. Meissner, DNA methylation in mammalian development and disease, Nat. Rev. Genet. 26 (2024) 7–30.

[2] K. M. Cherry, L. Qian, Supervised learning in DNA neural networks, Nature. 645 (2025) 639–647.

[3] Y.-R. Chang, J.-P. Shen, C.-F. Chou, DNA dynamics and organization in sub-micron scale: Bacterial chromosomes and plasmids in vivo and in vitro, Chin. J. Phys. 66 (2020) 82–90.

[4] A. R. Chandrasekaran, Nuclease resistance of DNA nanostructures, Nat. Rev. Chem. 5 (2021) 225–239.

[5] D. R. McCarthy, J. M. Remington, J. B. Ferrell, S. T. Schneebeli, J. Li, Nano-Bio interactions between DNA nanocages and human serum albumin, J. Chem. Theory Comput. 19 (2023) 7873–7881.

[6] A. M. Mohammed, RETRACTED: Au Nanoparticle-Modified SiO2 Thin Film for DNA Immobilization and Hybridization, Chin. J. Phys. 56 (2018) 3099.

[7] C. E. Carr, L. A. Marky, Effect of GCAA stabilizing loops on three-and four-way intramolecular junctions, Phys. Chem. Chem. Phys. 20 (2018) 5046–5056.

[8] B. Akabayov, S. R. Akabayov, S.-J. Lee, G. Wagner, C. C. Richardson, Impact of macromolecular crowding on DNA replication, Nat. Commun. 4 (2013) 1615.

[9] Y. Sasaki, D. Miyoshi, N. Sugimoto, Effect of molecular crowding on DNA polymerase activity, Biotechnol. J.: Healthc. Nutr. Technol. 1 (2006) 440–446.

[10] S. B. Zimmerman, B. Harrison, Macromolecular crowding increases binding of DNA polymerase to DNA: an adaptive effect, Proc. Natl. Acad. Sci. U.S.A. 84 (1987) 1871–1875.

[11] D. Miyoshi, N. Sugimoto, Molecular crowding effects on structure and stability of DNA, Biochimie. 90 (2008) 1040–1051.

[12] D. Miyoshi, H. Karimata, N. Sugimoto, Hydration regulates the thermodynamic stability of DNA structures under molecular crowding conditions, Nucleosides Nucleotides Nucleic Acids. 26 (2007) 589–595.

[13] C. Feng, Y.-L. Tan, Y.-X. Cheng, Y.-Z. Shi, Z.-J. Tan, Salt-dependent RNA pseudoknot stability: effect of spatial confinement, Front. Mol. Biosci. 8 (2021) 666369.

[14] X. Wang, Y.-Z. Shi, 3D structure and stability prediction of DNA with multi-way junctions in ionic solutions, PLoS Comput. Biol. 21 (2025) e1013346.

[15] A. Singh, A. Maity, N. Singh, Structure and Dynamics of dsDNA in Cell-like Environments, Entropy. 24 (2022) 1587.

[16] F. Liu, Y. Yang, X. Wan, H. Gao, Y. Wang, J. Lu, L.-P. Xu, S. Wang, Space-confinment-enhanced fluorescence detection of DNA on hydrogel particles array, Acs Nano. 16 (2022) 6266–6273.

[17] A. Pruska, J. A. Harrison, A. Granzhan, A. Marchand, R. Zenobi, Solution and gas-phase stability of DNA junctions from temperature-controlled electrospray ionization and surface-induced dissociation, Anal. Chem. 95 (2023) 14384–14391.

[18] F. W. Starr, W. Wang, L. M. Nocka, B. Z. Wiemann, D. M. Hinckley, I. Mukerji, Holliday junction thermodynamics and structure: comparisons of coarse-grained simulations and experiments, Biophys. J. 110 (2016) 178a.

[19] J. Spitzer, B. Poolman, The role of biomacromolecular crowding, ionic strength, and physicochemical gradients in the complexities of life’s emergence, Microbiol. Mol. Biol. Rev. 73 (2009) 371–388.

[20] D. Collette, D. Dunlap, L. Finzi, Macromolecular crowding and DNA: Bridging the gap between in vitro and in vivo, Int. J. Mol. Sci. 24 (2023) 17502.

[21] K. S. Harve, R. Lareu, R. Rajagopalan, M. Raghunath, Understanding how the crowded interior of cells stabilizes DNA/DNA and DNA/RNA hybrids–in silico predictions and in vitro evidence, Nucleic Acids Res. 38 (2010) 172–181.

[22] K. M. Ravikumar, W. Huang, S. Yang, Fast-SAXS-pro: a unified approach to computing SAXS profiles of DNA, RNA, protein, and their complexes, Biophys. J. 104 (2013) 183a.

[23] R. P. Rambo, J. A. Tainer, Bridging the solution divide: comprehensive structural analyses of dynamic RNA, DNA, and protein assemblies by small-angle X-ray scattering, Curr. Opin. Struct. Biol. 20 (2010) 128–137.

[24] M. F. Ober, A. Baptist, L. Wassermann, A. Heuer-Jungemann, B. Nickel, In situ small-angle X-ray scattering reveals strong condensation of DNA origami during silicification, Nat. Commun. 13 (2022) 5668.

[25] A. K. Sieradzan, A. Gieldon, Y. Yin, Y. He, H. A. Scheraga, A. Liwo, A new protein nucleic-acid coarse -grained force field based on the UNRES and NARES-2P force fields, J. Comput. Chem. 39 (2018) 2360–2370.

[26] A. L. Forget, S. C. Kowalczykowski, Single-molecule imaging of DNA pairing by RecA reveals a three-dimensional homology search, Nature. 482 (2012) 423–427.

[27] J. Gorman, F. Wang, S. Redding, A. J. Plys, T. Fazio, S. Wind, E. E. Alani, E. C. Greene, Single-molecule imaging reveals target-search mechanisms during DNA mismatch repair, Proc. Natl. Acad. Sci. U.S.A. 109 (2012) E3074–E3083.

[28] M. Kümmerlin, Q. Zhao, J. Hazra, C. Hepp, A. Farrar, P. Turner, A. N. Kapanidis, Tunable fluorogenic DNA probes drive fast and high-resolution single-molecule fluorescence imaging, Nucleic Acids Res. 53 (2025) gkaf593.

[29] J. Abramson, J. Adler, J. Dunger, R. Evans, T. Green, A. Pritzel, O. Ronneberger, L. Willmore, A. J. Ballard, J. Bambrick, Accurate structure prediction of biomolecular interactions with AlphaFold 3, Nature. 630 (2024) 493–500.

[30] J. Jumper, R. Evans, A. Pritzel, T. Green, M. Figurnov, O. Ronneberger, K. Tunyasuvunakool, R. Bates, A. Žídek, A. Potapenko, Highly accurate protein structure prediction with AlphaFold, Nature. 596 (2021) 583–589.

[31] Y. Xiong, Y. Zhang, J. Wang, Y. Xiao, Using 3dRNA/DNA for RNA and DNA 3D Structure Prediction and Evaluation, Curr. Protoc. 3 (2023) e770.

[32] Y. Zhang, Y. Xiong, C. Yang, Y. Xiao, 3dRNA/DNA: 3D Structure Prediction from RNA to DNA, J. Mol. Biol. 436 (2024) 168742.

[33] S. Scott, C. Shaheen, B. McGuinness, K. Metera, F. Kouzine, D. Levens, C. J. Benham, S. Leslie, Single-molecule visualization of the effects of ionic strength and crowding on structure-mediated interactions in supercoiled DNA molecules, Nucleic Acids Res. 47 (2019) 6360–6368.

[34] W. He, X. Qiu, S. Kirmizialtin, Sequence-dependent orientational coupling and electrostatic attraction in cation-mediated DNA–DNA interactions, J. Chem. Theory Comput. 19 (2023) 6827–6838.

[35] D. M. Hinckley, G. S. Freeman, J. K. Whitmer, J. J. De Pablo, An experimentally-informed coarse-grained 3-site-per-nucleotide model of DNA: Structure, thermodynamics, and dynamics of hybridization, J. Chem. Phys. 139 (2013) 144903.

[36] G. S. Freeman, D. M. Hinckley, J. P. Lequieu, J. K. Whitmer, J. J. De Pablo, Coarse-grained modeling of DNA curvature, J. Chem. Phys. 141 (2014) 165103.

[37] T. E. Ouldridge, A. A. Louis, J. P. Doye, Structural, mechanical, and thermodynamic properties of a coarse-grained DNA model, J. Chem. Phys. 134 (2011) 085101.

[38] G. S. Freeman, D. M. Hinckley, J. J. de Pablo, A coarse-grain three-site-per-nucleotide model for DNA with explicit ions, J. Chem. Phys. 135 (2011)

[39] E. Poppleton, R. Romero, A. Mallya, L. Rovigatti, P. Šulc, OxDNA. org: a public webserver for coarse-grained simulations of DNA and RNA nanostructures, Nucleic Acids Res. 49 (2021) W491–W498.

[40] F. Hong, J. S. Schreck, P. Šulc, Understanding DNA interactions in crowded environments with a coarse-grained model, Nucleic Acids Res. 48 (2020) 10726–10738.

[41] H. Chhabra, G. Mishra, Y. Cao, D. Presern, E. Skoruppa, M. M. Tortora, J. P. Doye, Computing the elastic mechanical properties of rodlike DNA nanostructures, J. Chem. Theory Comput. 16 (2020) 7748–7763.

[42] A. K. Sieradzan, Ł. Golon, A. Liwo, Prediction of DNA and RNA structure with the NARES-2P force field and conformational space annealing, Phys. Chem. Chem. Phys. 20 (2018) 19656–19663.

[43] S. Pasquali, P. Derreumaux, HiRE-RNA: a high resolution coarse-grained energy model for RNA, J. Phys. Chem. B. 114 (2010) 11957–11966.

[44] D. Chakraborty, N. Hori, D. Thirumalai, Sequence-dependent three interaction site model for single-and double-stranded DNA, J. Chem. Theory Comput. 14 (2018) 3763–3779.

[45] J. J. Uusitalo, H. I. Ingólfsson, P. Akhshi, D. P. Tieleman, S. J. Marrink, Martini coarse-grained force field: extension to DNA, J. Chem. Theory Comput. 11 (2015) 3932–3945.

[46] A. K. Sieradzan, C. R. Czaplewski, E. A. Lubecka, A. G. Lipska, A. S. Karczynska, A. P. Gieldon, R. Slusarz, M. Makowski, P. Krupa, M. Kogut, Extension of the Unres package for physics-based coarse-grained simulations of proteins and protein complexes to very large systems, Biophys. J. 120 (2021) 83a–84a.

[47] M. Ullner, C. E. Woodward, B. Jönsson, A Debye–Hückel theory for electrostatic interactions in proteins, J. Chem. Phys. 105 (1996) 2056–2065.

[48] Z.-C. Mu, Y.-L. Tan, J. Liu, B.-G. Zhang, Y.-Z. Shi, Computational modeling of DNA 3D structures: From dynamics and mechanics to folding, Molecules. 28 (2023) 4833.

[49] O. Coskuner-Weber, M. Koca, V. N. Uversky, Molecular Crowding by Computational Approaches, (Macro) Molecular Crowding: Life of the Pottage. (2025) 471–497.

[50] Z.-C. Mu, Y.-L. Tan, B.-G. Zhang, J. Liu, Y.-Z. Shi, Ab initio predictions for 3D structure and stability of single-and double-stranded DNAs in ion solutions, PLoS Comput. Biol. 18 (2022) e1010501.

[51] G. S. Manning, The molecular theory of polyelectrolyte solutions with applications to the electrostatic properties of polynucleotides, Q. Rev. Biophys. 11 (1978) 179–246.

[52] Z.-J. Tan, S.-J. Chen, Electrostatic free energy landscapes for nucleic acid helix assembly, Nucleic Acids Res. 34 (2006) 6629–6639.

[53] Z. J. Tan, S. J. Chen, RNA helix stability in mixed Na+/Mg2+ solution, Biophys. J. 92 (2007) 3615–3632.

[54] X. Wang, Y. L. Tan, S. Yu, Y. Z. Shi, Z. J. Tan, Predicting 3D structures and stabilities for complex RNA pseudoknots in ion solutions, Biophys. J. 122 (2023) 1503–1516.

[55] M. Feig, I. Yu, P.-h. Wang, G. Nawrocki, Y. Sugita, Crowding in cellular environments at an atomistic level from computer simulations, J. Phys. Chem. B. 121 (2017) 8009–8025.

[56] N. F. Dupuis, E. D. Holmstrom, D. J. Nesbitt, Molecular-crowding effects on single-molecule RNA folding/unfolding thermodynamics and kinetics, Proc. Natl. Acad. Sci. U.S.A. 111 (2014) 8464–8469.

[57] M. S. Cheung, D. Klimov, D. Thirumalai, Molecular crowding enhances native state stability and refolding rates of globular proteins, Proc. Natl. Acad. Sci. U.S.A. 102 (2005) 4753–4758.

[58] S. Qin, H.-X. Zhou, Further development of the FFT-based method for atomistic modeling of protein folding and binding under crowding: Optimization of accuracy and speed, J. Chem. Theory Comput. 10 (2014) 2824–2835.

[59] J. Mittal, R. B. Best, Thermodynamics and kinetics of protein folding under confinement, Proc. Natl. Acad. Sci. U.S.A. 105 (2008) 20233–20238.

[60] D. Klimov, D. Newfield, D. Thirumalai, Simulations of β-hairpin folding confined to spherical pores using distributed computing, Proc. Natl. Acad. Sci. U.S.A. 99 (2002) 8019–8024.

[61] Z.-J. Tan, S.-J. Chen, Ion-mediated RNA structural collapse: effect of spatial confinement, Biophys. J. 103 (2012) 827–836.

[62] K. Hukushima, K. Nemoto, Exchange Monte Carlo method and application to spin glass simulations, J. Phys. Soc. Japan. 65 (1996) 1604–1608.

[63] D. Gront, A. Kolinski, J. Skolnick, A new combination of replica exchange Monte Carlo and histogram analysis for protein folding and thermodynamics, J. Chem. Phys. 115 (2001) 1569–1574.

[64] Y. Z. Shi, F. H. Wang, Y. Y. Wu, Z. J. Tan, A coarse-grained model with implicit salt for RNAs: predicting 3D structure, stability and salt effect, J. Chem. Phys. 141 (2014) 105102.

[65] Y. Okamoto, Generalized-ensemble algorithms: enhanced sampling techniques for Monte Carlo and molecular dynamics simulations, J. Mol. Graph. Model. 22 (2004) 425–439.

[66] H. Fukunishi, O. Watanabe, S. Takada, On the Hamiltonian replica exchange method for efficient sampling of biomolecular systems: Application to protein structure prediction, J. Chem. Phys. 116 (2002) 9058–9067.

[67] J. D. Chodera, W. C. Swope, J. W. Pitera, C. Seok, K. A. Dill, Use of the weighted histogram analysis method for the analysis of simulated and parallel tempering simulations, J. Chem. Theory Comput. 3 (2007) 26–41.

[68] D. Bouzida, P. A. Rejto, G. M. Verkhivker, Monte Carlo study of ligand–protein binding energy landscapes with the weighted histogram analysis method, Int. J. Quantum Chem. 73 (1999) 113–121.

[69] E. Gallicchio, M. Andrec, A. K. Felts, R. M. Levy, Temperature weighted histogram analysis method, replica exchange, and transition paths, J. Phys. Chem. B. 109 (2005) 6722–6731.

[70] M. J. Boniecki, G. Lach, W. K. Dawson, K. Tomala, P. Lukasz, T. Soltysinski, K. M. Rother, J. M. Bujnicki, SimRNA: a coarse-grained method for RNA folding simulations and 3D structure prediction, Nucleic Acids Res. 44 (2016) e63–e63.

[71] J. Stasiewicz, S. Mukherjee, C. Nithin, J. M. Bujnicki, QRNAS: software tool for refinement of nucleic acid structures, BMC Struct. Biol. 19 (2019) 5.

[72] T. Kanungo, D. M. Mount, N. S. Netanyahu, C. Piatko, R. Silverman, A. Y. Wu, in Proceedings of the sixteenth annual symposium on Computational geometry. (2000), pp. 100–109.

[73] Y. Z. Shi, L. Jin, C. J. Feng, Y. L. Tan, Z. J. Tan, Predicting 3D structure and stability of RNA pseudoknots in monovalent and divalent ion solutions, PLoS Comput. Biol. 14 (2018) e1006222.

[74] Y. Zhao, Y. Huang, Z. Gong, Y. Wang, J. Man, Y. Xiao, Automated and fast building of three-dimensional RNA structures, Sci. Rep. 2 (2012) 734.

[75] D. Zhang, S. J. Chen, IsRNA: An iterative simulated reference state approach to modeling correlated interactions in RNA folding, J. Chem. Theory Comput. 14 (2018) 2230–2239.

[76] J. Singh, J. Hanson, K. Paliwal, Y. Zhou, RNA secondary structure prediction using an ensemble of two-dimensional deep neural networks and transfer learning, Nat. Commun. 10 (2019) 5407.

[77] J. A. Cruz, M. F. Blanchet, M. Boniecki, J. M. Bujnicki, S. J. Chen, S. Cao, R. Das, F. Ding, N. V. Dokholyan, S. C. Flores, RNA-Puzzles: a CASP-like evaluation of RNA three-dimensional structure prediction, RNA. 18 (2012) 610–625.

[78] R. Blake, S. G. Delcourt, Thermal stability of DNA, Nucleic Acids Res. 26 (1998) 3323–3332.

[79] M.-H. Hou, S.-B. Lin, J.-M. P. Yuann, W.-C. Lin, A. H.-J. Wang, L.-s. Kan, Effects of polyamines on the thermal stability and formation kinetics of DNA duplexes with abnormal structure, Nucleic Acids Res. 29 (2001) 5121–5128.

[80] J. Lah, S. Hadzi, Thermodynamic Origin of the Linear Pressure Dependence of DNA Thermal Stability, J. Phys. Chem. Lett. 15 (2024) 9064–9069.

[81] Z.-Y. Song, X. Zhang, X. Ai, L.-Y. Huang, X.-M. Hou, P. Fossé, N.-N. Liu, O. Mauffret, S. Réty, X.-G. Xi, Structural mechanism of RECQ1 helicase in unfolding G-quadruplexes compared with duplex DNA, Nucleic Acids Res. 53 (2025) gkaf877.

[82] R. Smith, T. Lebeaupin, S. Juhász, C. Chapuis, O. D’Augustin, S. Dutertre, P. Burkovics, C. Biertümpfel, G. Timinszky, S. Huet, Poly (ADP-ribose)-dependent chromatin unfolding facilitates the association of DNA-binding proteins with DNA at sites of damage, Nucleic Acids Res. 47 (2019) 11250–11267.

[83] A. R. Gruber, R. Lorenz, S. H. Bernhart, R. Neuböck, I. L. Hofacker, The vienna RNA websuite, Nucleic Acids Res. 36 (2008) W70–W74.

[84] W. Humphrey, A. Dalke, K. Schulten, VMD: visual molecular dynamics, J. Mol. Graph. 14 (1996) 33–38.

