## Supplementary materials for "Spatial confinement reshapes the folding of an ion-stabilized DNA with three-way junction"

#### The coarse-grained force filed of our DNAfold2 model

The total potential energy $U$ in our DNAfold2 model comprises eight distinct components, as detailed in [1].

$$U = U_b + U_a + U_d + U_{bs} + U_{bp} + U_{exc} + U_{cs} + U_{el} + U_{conf}, \tag{S1}$$

the first three terms correspond to bonded energy contributions: $U_b$ for virtual bond lengths, $U_a$ for bond angles, and $U_d$ for dihedral angles, collectively describing the connectivity and local geometry of the DNA chain. The explicit forms of $U_b$, $U_a$, and $U_d$ are provided in [2].

$$U_b = \sum_{bonds} K_b (r - r_0)^2; \tag{S2}$$

$$U_a = \sum_{angles} K_\theta (\theta - \theta_0)^2; \tag{S3}$$

$$U_d = \sum_{dihedrals} \left\{ K_\varphi [1 - \cos(\varphi - \varphi_0)] + \frac{1}{2} K_\varphi [1 - \cos 3(\varphi - \varphi_0)] \right\}, \tag{S4}$$

the parameters $K_b$, $K_\theta$, and $K_\varphi$ represent the energy strengths, while $r_0$, $\theta_0$, and $\varphi_0$ denote the equilibrium distances and angles for virtual bonds, bond angles, and dihedral angles, respectively, at the energy minimum. To determine these bonded energy parameters, we followed a systematic procedure. First, a statistical analysis of 138 experimental DNA structures from the Protein Data Bank (including both ssDNA and dsDNA; see Table S1) was performed to obtain the distance and angle distributions of the coarse-grained atoms. Next, the functional forms in Eqs. S2–S5 were fitted to these distributions, from which the bonded energy parameters were derived using the following equation

$$U(x) = -k_B T \ln[P(x)], \tag{S5}$$

here, $k_B$ is the Boltzmann constant and $T$ the absolute temperature in Kelvin. $P_x$ denotes the normalized distributions of the bonded distances and angles $x$. The model defines two sets of bonded energy parameters: $Para_{loop}$ and $Para_{helix}$. The $Para_{loop}$ energy models DNA folding as a free chain and is derived from single-stranded or loop regions (non-helix parts) in experimental structures. In contrast, the $Para_{helix}$ energy is applied during structure refinement, derived from stem (helix) regions in experimental structures, and also used for base-pairing nucleotides in native-like structures predicted during folding. Further details are provided in Refs [1].

In Eq. S1, $U_{bs}$ represents the base-stacking interaction between adjacent base pairs. Its functional form is given by:

$$U_{bs} = \frac{1}{2} \sum_{i,j}^{N_{st}} \left| G_{i,i+1,j-1,j} \right| \left\{ \left[ 5\left(\frac{\sigma_{st}}{r_{i,i+1}}\right)^{12} - 6\left(\frac{\sigma_{st}}{r_{i,i+1}}\right)^{10} \right] + \left[ 5\left(\frac{\sigma_{st}}{r_{j,j+1}}\right)^{12} - 6\left(\frac{\sigma_{st}}{r_{j,j+1}}\right)^{10} \right] \right\}, \tag{S6}$$

here, $\sigma_{st}$ denotes the optimal distance between two neighboring bases in known helical structures. $G_{i,i+1,j-1,j}$ represents the base-stacking energy strength, estimated by combining experimental thermodynamic parameters with Monte Carlo simulations based on our previous model [2,3]:

$$G_{i,i+1,j-1,j} = \Delta H - T(\Delta S - \Delta S_C). \quad \text{(S7)}$$

In the DNAfold2 model, $\Delta H$ and $\Delta S$ denote the DNA thermodynamic parameters derived from experimental studies [4,5]. Here, $\Delta S_C$ denotes the conformational entropy change, inherently captured by the Monte Carlo (MC) algorithm due to base-pair stacking. $\Delta S_C$ was computed via MC simulations of the DNA helix (Fig. S5), where the DNA molecule was fixed except for nucleotides with indices $i \leq$ or $\geq j$. The number of conformations $\Omega$ satisfying the stacking condition between pairs $(i, j)$ and $(i+1, j-1)$ was counted without imposing additional base-pairing or stacking constraints. The conformational entropy change for base stacking between these pairs was then calculated by:

$$\Delta S_c = k_B \ln(\Omega/\Omega_0), \quad \text{(S8)}$$

here, $k_B$ is the Boltzmann constant, and $\Omega_0$ denotes the total number of conformations sampled during the simulation. $\Delta S_c$ shows minimal variation across different base-pair positions iii; therefore, an average value of –11.5 eu was adopted in the present model for simplicity.

The base-pairing potential, calculated for all possible pairs (G–C, G–U, and A–U), is expressed as:

$$U_{bp} = \sum_{i<j-3}^{N_{bp}} \frac{\varepsilon_{bp}}{1+k_{NN}(r_{N_iN_j}-r_{NN})^2+k_{CN}\sum_{i(j)}\left(r_{C_iN_j}-r_{CN}\right)^2+k_{PN}\sum_{i(j)}\left(r_{P_iN_j}-r_{PN}\right)^2}, \quad \text{(S9)}$$

where $\varepsilon_{bp}$ is the interaction strength $\varepsilon_{AT}$ =2*$\varepsilon_{GC}$/3. $r_{NN}$, $r_{CN}$, and $r_{PN}$ are three distances between the corresponding atoms of P, C and N in two paired nucleotides to describe the orientation of hydrogen-bonding interactions, and the values of them were obtained from the pairing bases in the PDB structures. Additionally, $k_{NN}$, $k_{CN}$ and $k_{PN}$ in Eq. S6 are the corresponding energy strength.

In Eq. S1, $U_{exc}$ represents the repulsive volume interaction between CG beads, modeled using a purely repulsive Lennard-Jones potential.

$$U_{exc} = \sum_{i<j}^{N} \begin{cases} 4\varepsilon\left[\left(\frac{\sigma_0}{r_{ij}}\right)^{12} - \left(\frac{\sigma_0}{r_{ij}}\right)^{6}\right], & if\ r_{ij} < \sigma_0 \\ 0, & if\ r_{ij} \geq \sigma_0 \end{cases}, \quad \text{(S10)}$$

here, $\varepsilon = 0.26$ kcal/mol represents the interaction strength, $\sigma_0$ is the sum of the radii of beads $i$ and $j$, and $r_{ij}$ is the distance between beads $i$ and $j$.

In Eq. S1, $U_{cs}$ denotes the coaxial-stacking interaction between adjacent base pairs from two discontinuous stems [1,6].

$$U_{cs} = \frac{1}{2} \sum_{i-j,k-l}^{N_{cst}} |G_{i,k,l,j}| \left\{ \left[1 - e^{-a(r_{ik}-r_{cs})}\right]^2 - \left[1 - e^{-a(r_{jl}-r_{cs})}\right]^2 - 2 \right\}, \quad \text{(S11)}$$

here, $G_{i,k,l,j}$ denotes the base-stacking energy strength, approximating interactions between discontinuous stems and their nearest-neighbor base pairs. The distances $r_{ik}$ ($r_{jl}$) correspond to those between interfacing bases $i(j)$ and $k(l)$ of the two stems. The parameter $a$ defines the range of the coaxial-stacking interaction, and $r_{cs}$ is the optimal distance between coaxially stacked stems, determined from statistical analysis of known PDB structures.

$U_{el}$ in Eq. S1 represents the electrostatic interaction between phosphates, with their reduced charges determined by the counterion condensation model [7] and the tightly bound ion model [8-10]

$$U_{el} = \sum_{i<j}^{N} \frac{Q_i Q_j e^2}{4\pi\epsilon_0 \epsilon r_{ij}} e^{-\frac{r_{ij}}{l_D}}, \quad \text{(S12)}$$

where $r_{ij}$ represents the distance between the $i$-th and $j$-th phosphate beads, and $l_D$ is the Debye length. The reduced charge on the $i$-th phosphate bead is

$$Q_i = 1 - f_i, \quad \text{(S13)}$$

where $f_i$ represents the ion neutralization fraction for the $i$-th phosphate bead. In addition to the assumption of a uniform distribution of binding ions along the DNA strand, $f_i$ depends on the DNA structure and incorporates both monovalent and divalent ions

$$f_i = x f_i^1 + (1-x) f_i^2, \quad \text{(S14)}$$

where $f_i^v$ $(v = 1, 2)$ represents the binding fraction of $v$-valent ions for the $i$-th phosphate bead. The terms $x$ and $(1-x)$ denote the contribution fractions of monovalent and divalent ions, respectively. In Eq. S13, which is derived from the TBI model [8-10]. When using $Na^+$ and $Mg^{2+}$ to represent monovalent and divalent ions, $x$ can be expressed using an empirical formula [8-10].

$$x = \frac{[\mathrm{Na}^+]}{[\mathrm{Na}^+] + \alpha[\mathrm{Mg}^{2+}]}, \quad \text{(S15)}$$

where $\alpha = (8.1 - 64.8/N)(5.2 - \ln[\mathrm{Na}^+])$, $[Na^+]$ and $[Mg^{2+}]$ represent the bulk concentrations of sodium and magnesium ions, respectively, and $N$ is the length of the DNA chain [8-10]. For further details, please refer to Refs [8-11].

To achieve a more precise electrostatic potential for an DNA structure, $f_i^v$ is defined as

$$f_i^v = \frac{N \bar{f}_i^v}{\sum_N e^{-\beta v \phi_i}} e^{-\beta v \phi_i}, \quad \text{(S16)}$$

where $\bar{f}_i^\nu = 1 - (\frac{b}{\nu l_B})$ represents the average neutralization fraction for the $i$-th phosphate bead [5]. In this expression, $b$ denotes the average charge spacing along the DNA backbone, and $l_B$ is the Bjerrum length. The electrostatic potential $\phi_i$ for the $i$-th phosphate bead can be approximated as follows:

$$\phi_i = \sum_{i \neq j}^{N} \frac{l_B Q_j}{r_{ij}} e^{-\frac{r_{ij}}{l_D}}. \tag{S17}$$

The reduced fraction $Q_i$ based on the DNA structure is determined through an iterative procedure: (1) Start by calculating the neutralization fraction $f_i^\nu$ using Eq. S14, with the initial value $f_i^\nu = \bar{f}_i^\nu$; (2) Compute $Q_i$ using Eq. S13, then substitute $Q_i$ into Eq. S17 to determine $\phi_i$; (3) Use Eq. S16 to calculate the fraction $f_i^\nu$ for the $\nu$-valent ion; (4) Repeat steps (1) through (3) until the value of $f_i^\nu$ converges.

**Weighted histogram analysis method**

In our CG model, we employed the Weighted Histogram Analysis Method (WHAM) [12] to calculate the fractions of each DNA state at various temperatures, utilizing REMC trajectories. This approach enables the quantitative analysis of DNA thermal stability, encompassing key properties such as melting temperatures and thermally induced unfolding pathways. Specifically, the thermal stability of a DNA molecule is evaluated through the following four steps. First, replica exchange Monte Carlo (REMC) simulations are performed across a range of temperatures to generate equilibrium trajectories. Second, the relevant reaction coordinates-namely, the structural states ($S$) and conformational energies ($E$)-are discretized into bins: $S_j$ ($j$=1, 2, ..., F) and $E_k$ ($k$=1, 2, ...,100). Here, $S$ represents distinct structural states (e.g., folded, unfolded, and intermediate), while $E$ corresponds to the energy levels of sampled conformations. Each pair of indices $(i, j)$ thus defines a microstate within the WHAM framework. As an illustrative example, we consider a DNA molecule with a three-way junction as a representative structural state. The probability of microstate $(j, k)$ at the $i$-th temperature, denoted $p_{i,(j,k)}$, is given by the following equation:

$$p_{i,(j,k)} = Z_i c_{i(j,k)} \mathring{p}_{(j,k)}, \tag{S18}$$

where $\mathring{p}_{(j,k)}$ represents the unbiased probability of a small state $(j, k)$ at the temperature of interest, with the partition function $Z_i$ chosen such that the sum of all small states satisfies $\sum_{j,k} p_{i,(j,k)} = 1$, ensuring normalization. The temperature-biasing factor $c_{i,(j,k)}$ is defined as $c_{i,(j,k)} = \exp[-(\beta_i - \beta_0)E_k]$, where $\beta_i$ is the inverse temperature at the *i*-th temperature, and $E_k$ is the energy of conformation $k$.

The unbiased probabilities $\overset{\circ}{p}_{(j,k)}$ for each small state are then determined by iterating the relevant equations until convergence is achieved.

$$\overset{\circ}{p}_{(j,k)} = \frac{\sum_{i=1}^{M} n_{i,(j,k)}}{\sum_{i=1}^{M} N_i Z_i c_{i,(j,k)}}, \tag{S19}$$

$$Z_i^{-1} = \sum_{j,k} c_{i,(j,k)} \overset{\circ}{p}_{(j,k)}. \tag{S20}$$

where $M$ represents the number of replicas, with $M = 10$ in our case. The quantity $n_{i,(j,k)}$ denotes the count of small state $(j,k)$ at the *i*-th temperature, and $N_i$ is the total number of conformations at the *i*-th temperature. Initially, the partition function $Z_i$ for each temperature is set to 1. Finally, the fraction $f_{S_j}(T)$ of each structural state of DNA at temperature $T$ is calculated using the following equation

$$f_{S_j}(T) = \sum_{k=1}^{100} \overset{\circ}{p}_{(j,k)}, \tag{S21}$$

where $S_j$ represents the different structural states of DNA, including folded (F), unfolded (U), and intermediate (I) states. The fraction $f_{S_j}(T)$ can be used to investigate the thermal stability and unfolding pathways of DNAs. Specifically, the melting temperature is determined by fitting the fractions of the folded state $f_F(T)$ $(F = S_F)$ and the unfolded state $f_U(T)$ $(U = S_1)$ to a two-state model.

$$f_F(T) = \frac{1}{1+e^{(T-T_{m1})/dT_1}}; \tag{S22}$$

$$f_U(T) = 1 - \frac{1}{1+e^{(T-T_{m2})/dT_2}}. \tag{S23}$$

Here, $T_{m1}$ and $T_{m2}$ represent the melting temperatures corresponding to the folded-to-intermediate (F$\rightarrow I$) and intermediate-to-unfolded (I$\rightarrow$U) transitions, respectively. $dT_1$ and $dT_2$ are adjustable parameters that characterize the widths or sharpness of these transitions.

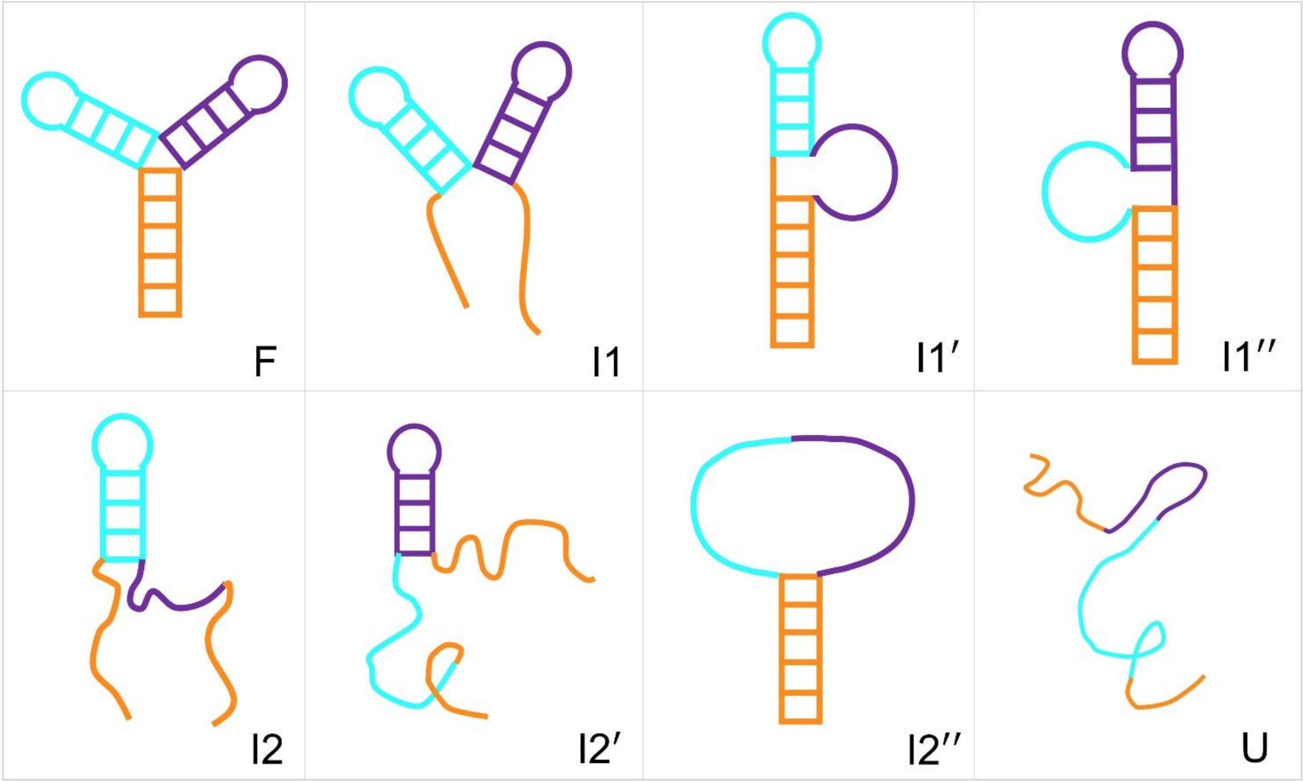


**Fig. S1.** Eight structural states of the 3WJ: one folded state (F), six intermediate states (I1, I1', I1'', I2, I2', I2''), and one unfolded state (U). Specifically, F: all three stems retained; I1: Stem 1 resolved; I1': Stem 3 resolved; I1'': Stem 2 resolved; I2: Stems 1 and 3 resolved; I2': Stems 1 and 2 resolved; I2'': Stems 2 and 3 resolved; U: all stems resolved.

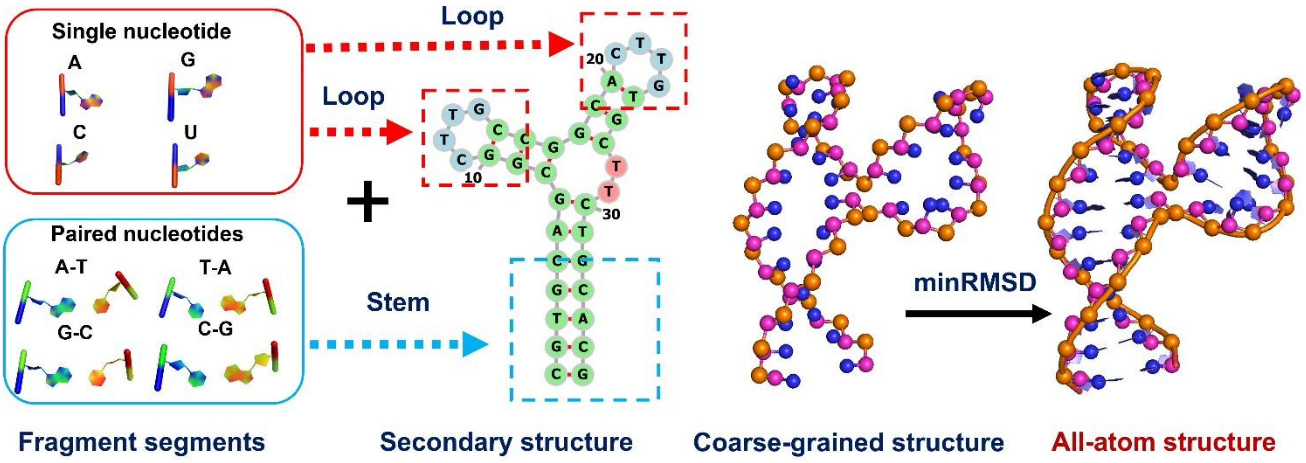


**Fig. S2.** Schematic illustration of the reconstruction of coarse-grained DNA structures into all-atom representations in our DNAfold2 model.

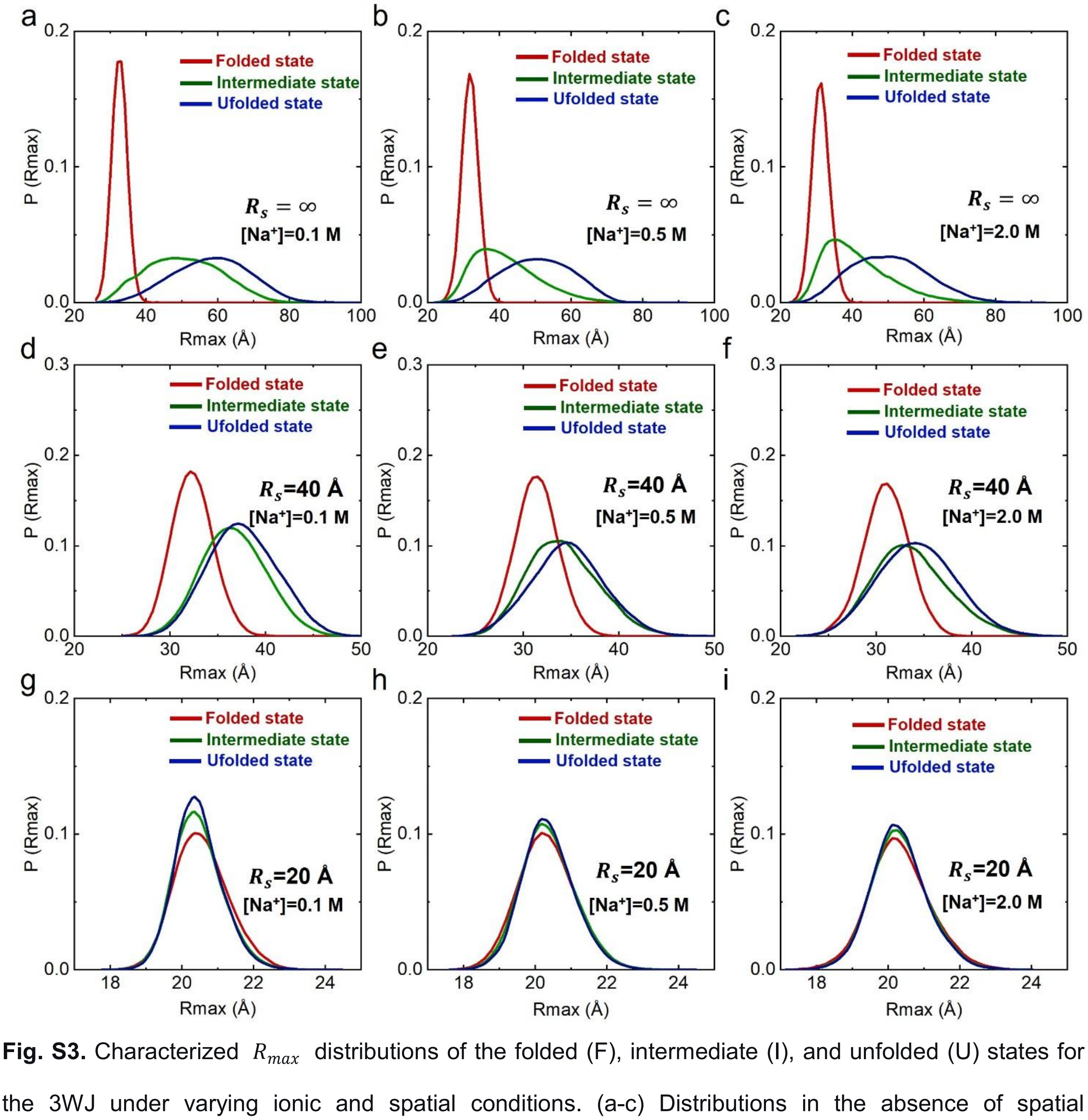


**Fig. S3.** Characterized $R_{max}$ distributions of the folded (F), intermediate (I), and unfolded (U) states for the 3WJ under varying ionic and spatial conditions. (a-c) Distributions in the absence of spatial confinement at [$Na^+$] of 0.1 M (a), 0.5 M (b), and 2 M (c). (d-f) Distributions under a spatial confinement of $R_s$ = 40 Å at 0.1 M (d), 0.5 M (e), and 2 M (f) $Na^+$. (g-i) Distributions under a spatial confinement of $R_s$ = 20 Å at 0.1 M (g), 0.5 M (h), and 2 M (i) $Na^+$.

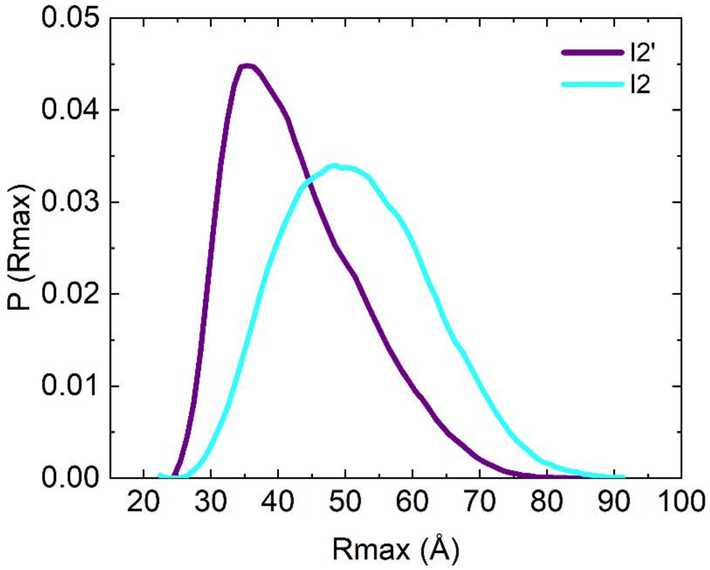


**Fig. S4.** Distributions of $R_{max}$ for the I2 and I2' states of the 3WJ under 1 M $Na^+$ in the absence of confinement.

**Table S1.** PDB codes of the 138 DNA structures used in our statistical analysis for constructing the coarse-grained (CG) force field.

| | | | | | | | | | |
|---|---|---|---|---|---|---|---|---|---|
| 1ag5 | 1agk | 1aw4 | 1bdz | 1b6x | 1cvx | 1cs7 | 1dxa | 1dnm | 1dcr |
| 1d13 | 1d16 | 1d49 | 1d63 | 1d89 | 1db6 | 1eek | 1ezn | 1en1 | 1en3 |
| 1fyk | 1fv8 | 1g6d | 1i0f | 1juu | 1lai | 1l0r | 1la8 | 1noq | 1puy |
| 1qdk | 1qph | 1qe7 | 1snj | 1sk5 | 1wqy | 1zfb | 1zfe | 1zfg | 1zfh |
| 1zyf | 1zyg | 1zew | 1zf9 | 107d | 116d | 119d | 126d | 158d | 183d |
| 195d | 196d | 2arg | 2b1b | 2b1d | 2d47 | 2f1q | 2gyx | 2kuz | 2k0v |
| 2k67 | 2k68 | 2k69 | 2lbi | 2lgm | 2lzv | 2lzw | 2lsc | 2ll9 | 2l13 |
| 2mav | 2mci | 2miv | 2mjj | 2mnf | 2m2c | 2npw | 2neo | 2org | 2pik |
| 2rrr | 2rvp | 2rt8 | 238d | 240d | 260d | 272d | 285d | 287d | 3co3 |
| 3gsj | 3l1q | 3omj | 3qsc | 3qk4 | 3r86 | 3v06 | 307d | 339d | 348d |
| 363d | 4e7y | 4f8g | 4j2i | 4kbd | 414d | 424d | 440d | 5ewb | 5gun |
| 5ip8 | 5ju4 | 5j3g | 5ki4 | 5mvp | 5mvq | 5m68 | 5uzf | 5xuv | 6asf |
| 6ast | 6dm7 | 6dy5 | 6g8s | 6iyq | 6ror | 6rou | 6s7d | 7b4z | 7edw |
| 7kcl | 7vck | 7ril | 7sb8 | 1aul | 1bdn | llp7 | 1bub | | |

**Table S2.** The parameters of bonded potentials of CG force field.

| Bond $U_b$ | | | | |
|---|---|---|---|---|
| | $K_b$ (kcal/mol/Å$^2$) | | $r_0$ (Å) | |
| | Helix | Loop | Helix | Loop |
| | $Para_{helix}$ | $Para_{loop}$ | $Para_{helix}$ | $Para_{loop}$ |
| $P_iC_i$ | 196.4 | 98.2 | 3.95 | 3.95 |
| $C_iP_{i+1}$ | 141.0 | 70.5 | 3.95 | 3.95 |
| $C_iN_i$ | 91.6 | 45.8 | 3.55 | 3.55 |
| **Angle $U_a$** | | | | |
| | $K_\theta$ (kcal/mol/rad$^2$) | | $\theta_0$ (rad) | |
| | Helix | Nonhelix | Helix | Nonhelix |
| | $Para_{helix}$ | $Para_{loop}$ | $Para_{helix}$ | $Para_{loop}$ |
| $P_iC_iP_{i+1}$ | 19.6 | 9.8 | 2.1 | 2.1 |
| $C_{i-1}P_iC_i$ | 17.2 | 8.6 | 1.8 | 1.8 |
| $P_iC_iN_{i+1}$ | 13.0 | 6.5 | 1.7 | 1.7 |
| $N_iC_iP_{i+1}$ | 28.6 | 14.3 | 1.7 | 1.7 |
| **Dihedral $U_d$** | | | | |
| | $K_\varphi$ (kcal/mol/rad$^2$) | | $\varphi_0$ (rad) | |
| | $Para_{helix}$ | $Para_{loop}$ | $Para_{helix}$ | $Para_{loop}$ |
| $P_iC_iP_{i+1}C_{i+1}$ | 2.6 | 1.3 | 2.5 | 2.5 |
| $C_{i-1}P_iC_iP_{i+1}$ | 8.0 | 4.0 | -2.9 | -2.9 |
| $C_{i-1}P_iC_iN_i$ | 6.4 | 3.2 | -1.3 | -1.3 |
| $N_{i-1}C_{i-1}P_iC_i$ | 3.2 | 1.6 | 0.9 | 0.9 |

The $Para_{helix}$ parameters are applied during the refinement of folded structures to describe base-pairing regions (stems), whereas the $Para_{loop}$ parameters are used in DNA folding simulations to represent flexible, unpaired regions and during the refinement of these regions.

**Table S3.** The parameters for the energy functions of base pairing and base stacking.

| Base pairing $U_{bp}$ | | | |
|---|---|---|---|
| Distance | $r_{NN}$ | $r_{CN}$ | $r_{PN}$ |
| $r$ (Å) | 8.9 | 12.1 | 14.1 |
| Energy strength | $K_{NN}$ | $K_{CN}$ | $K_{PN}$ |
| $k$ (kcal/mol) | 2.66 | 1.37 | 0.46 |
| $\varepsilon_{GC}$ =-3.3 kcal/mol; $\varepsilon_{AU}$ =-2.1 kcal/mol | | | |
| **Base stacking $U_{bs}$** | | | |
| $\Delta S_c$=-11.5 eu | | | |

**Table S4.** The predicted temperature of 3WJ with spatial confinement ( $R_s$ =20 Å) at extensive ion concentration by our present model.

| NO | Ion concentration (mM) | Melting temperature (℃) | |
|---|---|---|---|
| | [$Na^+$] | $T_{m1}{}^a$ | $T_{m2}{}^b$ |
| 1 | 10 | 45.4 | 90.3 |
| 2 | 50 | 47.4 | 95.8 |
| 3 | 100 | 50.1 | 105.4 |
| 4 | 200 | 52.2 | 110.1 |
| 5 | 500 | 54.6 | 115.5 |
| 6 | 1000 | 56.3 | 117.6 |
| 7 | 2000 | 56.8 | 118.3 |

$^aT_{m1}$ and $^bT_{m2}$ are the melting temperatures for the transitions from folded state to intermediate state and from intermediate state to unfolded state, respectively.

**Table S5.** The predicted temperature of 3WJ with spatial confinement ( $R_s$ =40 Å) at extensive ion concentration by our present model.

| NO | Ion concentration (mM) | Melting temperature (℃) | |
|---|---|---|---|
| | [$Na^+$] | $T_{m1}$[a] | $T_{m2}$[b] |
| 1 | 10 | 34.6 | 57.9 |
| 2 | 50 | 38.8 | 67.5 |
| 3 | 100 | 43 | 80 |
| 4 | 200 | 47.1 | 88.3 |
| 5 | 500 | 49 | 94.7 |
| 6 | 1000 | 49.8 | 97 |
| 7 | 2000 | 50.1 | 97.4 |

[a]$T_{m1}$ and [b]$T_{m2}$ are the melting temperatures for the transitions from folded state to intermediate state and from intermediate state to unfolded state, respectively.

**Table S6.** The predicted temperature of 3WJ without confinement at extensive ion concentration by our present model.

| NO | Ion concentration (mM) | Melting temperature (℃) | |
|---|---|---|---|
| | $[Na^+]$ | $T_{m1}{}^{a}$ | $T_{m2}{}^{b}$ |
| 1 | 10 | 30.8 | 52.5 |
| 2 | 50 | 32.4 | 55.7 |
| 3 | 100 | 34.4 | 64.6 |
| 4 | 200 | 38.7 | 70.6 |
| 5 | 500 | 42.9 | 75.3 |
| 6 | 1000 | 46.2 | 78.8 |
| 7 | 2000 | 47.2 | 79.2 |

${}^{a}T_{m1}$ and ${}^{b}T_{m2}$ are the melting temperatures for the transitions from folded state to intermediate state and from intermediate state to unfolded state, respectively.

### Reference


[1] Z.-C. Mu, Y.-L. Tan, B.-G. Zhang, J. Liu, Y.-Z. Shi, Ab initio predictions for 3D structure and stability of single-and double-stranded DNAs in ion solutions, *PLoS Comput. Biol*. 18 (2022) e1010501.

[2] X. Wang, Y. L. Tan, S. Yu, Y. Z. Shi, Z. J. Tan, Predicting 3D structures and stabilities for complex RNA pseudoknots in ion solutions, *Biophys. J*. 122 (2023) 1503-1516.

[3] Y. Z. Shi, F. H. Wang, Y. Y. Wu, Z. J. Tan, A coarse-grained model with implicit salt for RNAs: predicting 3D structure, stability and salt effect, *J. Chem. Phys*. 141 (2014) 105102.

[4] J. SantaLucia, H. T. Allawi, P. A. Seneviratne, Improved nearest-neighbor parameters for predicting DNA duplex stability, *Biochemistry*. 35 (1996) 3555-3562.

[5] J. SantaLucia Jr, D. Hicks, The thermodynamics of DNA structural motifs, *Annu. Rev. Biophys. Biomol. Struct.* 33 (2004) 415-440.

[6] Y. Z. Shi, L. Jin, C. J. Feng, Y. L. Tan, Z. J. Tan, Predicting 3D structure and stability of RNA pseudoknots in monovalent and divalent ion solutions, *PLoS Comput. Biol*. 14 (2018) e1006222.

[7] G. S. Manning, The molecular theory of polyelectrolyte solutions with applications to the electrostatic properties of polynucleotides, *Q. Rev. Biophys*. 11 (1978) 179-246.

[8] Z. J. Tan, S. J. Chen, Electrostatic correlations and fluctuations for ion binding to a finite length polyelectrolyte, *J. Chem. Phys*. 122 (2005) 044903.

[9] Z. J. Tan, S. J. Chen, Electrostatic free energy landscapes for nucleic acid helix assembly, *Nucleic Acids Res*. 34 (2006) 6629-6639.

[10] Z. J. Tan, S. J. Chen, Nucleic acid helix stability: effects of salt concentration, cation valence and size, and chain length, *Biophys. J*. 90 (2006) 1175-1190.

[11] L. Jin, Y. Z. Shi, C. J. Feng, Y. L. Tan, Z. J. Tan, Modeling structure, stability, and flexibility of double-stranded RNAs in salt solutions, *Biophys. J*. 115 (2018) 1403-1416.

[12] S. Kumar, J. M. Rosenberg, D. Bouzida, R. H. Swendsen, P. A. Kollman, The weighted histogram analysis method for free ‐energy calculations on biomolecules. I. The method, *J. Comput. Chem*. 13 (1992) 1011-1021.
